# A high-performance speech neuroprosthesis

**DOI:** 10.1101/2023.01.21.524489

**Authors:** Francis R. Willett, Erin M. Kunz, Chaofei Fan, Donald T. Avansino, Guy H. Wilson, Eun Young Choi, Foram Kamdar, Leigh R. Hochberg, Shaul Druckmann, Krishna V. Shenoy, Jaimie M. Henderson

**Author notes:** Co-first author (Francis R. Willett, Erin M. Kunz, Chaofei Fan). Co-senior author (Krishna V. Shenoy, Jaimie M. Henderson).

## Abstract

Speech brain-computer interfaces (BCIs) have the potential to restore rapid communication to people with paralysis by decoding neural activity evoked by attempted speaking movements into text^1,2^ or sound^3,4^. Early demonstrations, while promising, have not yet achieved accuracies high enough for communication of unconstrainted sentences from a large vocabulary^1–7^. Here, we demonstrate the first speech-to-text BCI that records spiking activity from intracortical microelectrode arrays. Enabled by these high-resolution recordings, our study participant, who can no longer speak intelligibly due amyotrophic lateral sclerosis (ALS), achieved a 9.1% word error rate on a 50 word vocabulary (2.7 times fewer errors than the prior state of the art speech BCI^2^) and a 23.8% word error rate on a 125,000 word vocabulary (the first successful demonstration of large-vocabulary decoding). Our BCI decoded speech at 62 words per minute, which is 3.4 times faster than the prior record for any kind of BCI^8^ and begins to approach the speed of natural conversation (160 words per minute^9^). Finally, we highlight two aspects of the neural code for speech that are encouraging for speech BCIs: spatially intermixed tuning to speech articulators that makes accurate decoding possible from only a small region of cortex, and a detailed articulatory representation of phonemes that persists years after paralysis. These results show a feasible path forward for using intracortical speech BCIs to restore rapid communication to people with paralysis who can no longer speak.

## Results

### Representation of speech production

It is not yet known how orofacial movement and speech production is organized in motor cortex at single neuron resolution. To investigate this, we recorded neural activity from four microelectrode arrays: two in area 6v (ventral premotor cortex)^10^ and two in area 44 (part of Broca’s area), while our study participant in the BrainGate2 pilot clinical trial attempted to make individual orofacial movements, speak single phonemes, or speak single words in response to cues displayed on a computer monitor (Fig. 1a-b; SFig 1 shows recorded spike waveforms). Implant locations for the arrays were chosen using the Human Connectome Project multi-modal cortical parcellation procedure^10^ (SFig 2). Our participant (T12) has bulbar-onset ALS and retains some limited orofacial movement and an ability to vocalize, but is unable to produce intelligible speech.

**Fig. 1.**
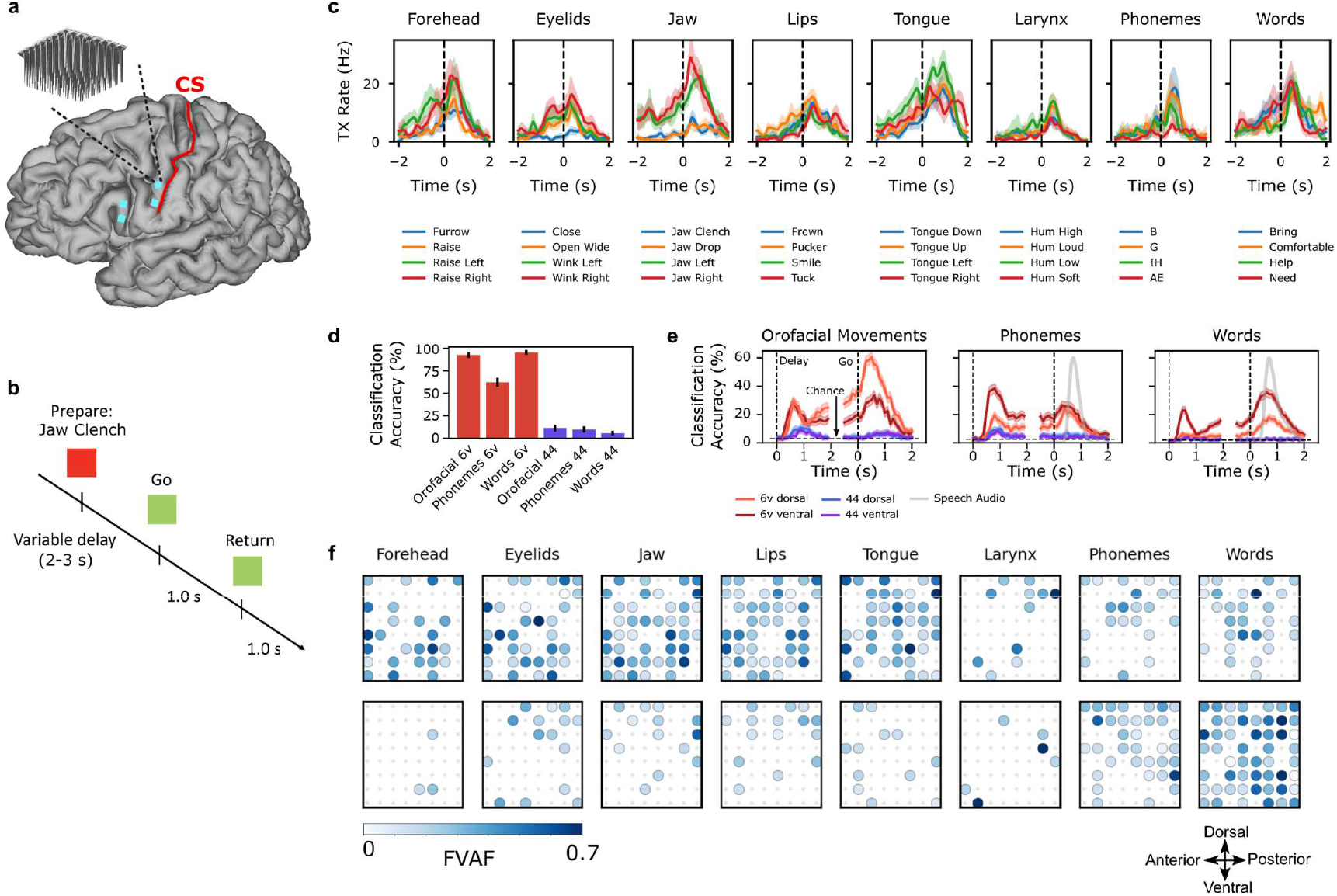
**(a)** Microelectrode array locations (cyan squares) are shown on top of MRI-derived brain anatomy (CS = central sulcus). **(b)** Neural tuning to orofacial movements, phonemes and words was evaluated in an instructed delay task. **(c)** Example responses of an electrode in area 6v that was tuned to a variety of speech articulator motions, phonemes and words. Each line shows the mean threshold crossing rate across all 16 trials of a single condition, and shaded regions show 95% CIs. Neural activity was denoised by convolving with a Gaussian smoothing kernel (80-ms SD). **(d)** Classification accuracy of a naïve Bayes decoder applied to 1 second of neural population activity from area 6v (red bars) or area 44 (purple bars) across all movement conditions (33 orofacial movements, 39 phonemes, 50 words). Black lines show 95% confidence intervals. **(e)** Classification accuracy across time for each of the four arrays and three types of movement. Classification was performed with a 100 ms window of neural population activity for each time point. Shaded regions show 95% confidence intervals. **(f)** Tuning heatmaps for both arrays in area 6v, for each movement category. Circles are drawn if binned firing rates on that electrode are significantly different across the given set of conditions (p<1e-5 assessed with a one-way ANOVA; bin width=800 ms). Shading indicates the fraction of variance accounted for by the across-condition differences in mean firing rate.

We found strong tuning to all tested categories of movement in area 6v (Fig 1c shows an example electrode). Neural activity in 6v was highly separable between movements: using a simple naive Bayes classifier applied to 1 second of neural population activity for each trial, we could decode from among 33 orofacial movements with 92% accuracy, 39 phonemes with 62% accuracy, and 50 words with 94% accuracy (Fig 1d; SFig 3). In contrast, although area 44 has previously been implicated in high-order aspects of speech production^11–14^, it appeared to contain little to no information about orofacial movements, phonemes or words (classification accuracy < 11%; Fig 1d). The absence of production-related neural activity in area 44 is consistent with some recent work questioning the traditional role of Broca’s area in speech^15–18^.

Next, we examined how information about each movement category was distributed across area 6v. We found that speech could be more accurately decoded from the ventral array, especially during the instructed delay period (Fig 1e), while the dorsal array contained more information about orofacial movements. This result is consistent with resting state fMRI data from the Human Connectome Project^10^ and from T12 that situates the ventral region of 6v as part of a language-related network (SFig 2). Nevertheless, both 6v arrays contained rich information about all movement categories. Finally, we found that tuning to speech articulators (jaw, larynx, lips, or tongue) was intermixed at the single electrode level (Fig 1f; SFig 4), and that all speech articulators were clearly represented within both 3.2 × 3.2 mm arrays. Although prior work using electrocorticographic grids has suggested that there may be a broader somatotopic organization^19^ along precentral gyrus, these results suggest that speech articulators are highly intermixed at a single neuron level.

In sum, robust and spatially intermixed tuning to all tested movements suggests that the representation of speech articulation is likely strong enough to support a speech BCI, despite paralysis and narrow coverage of the cortical surface. As area 44 appeared to contain little information about speech production, all further analyses were based on area 6v recordings only.

### Decoding attempted speech

Next, we tested whether we could neurally decode whole sentences in real-time. We trained a recurrent neural network (RNN) decoder to emit, at each 80 ms time step, the probability of each phoneme being spoken at that time. These probabilities were then combined with a language model to infer the most likely underlying sequence of words, given both the phoneme probabilities and the statistics of the English language (Fig 2a).

**Fig. 2.**
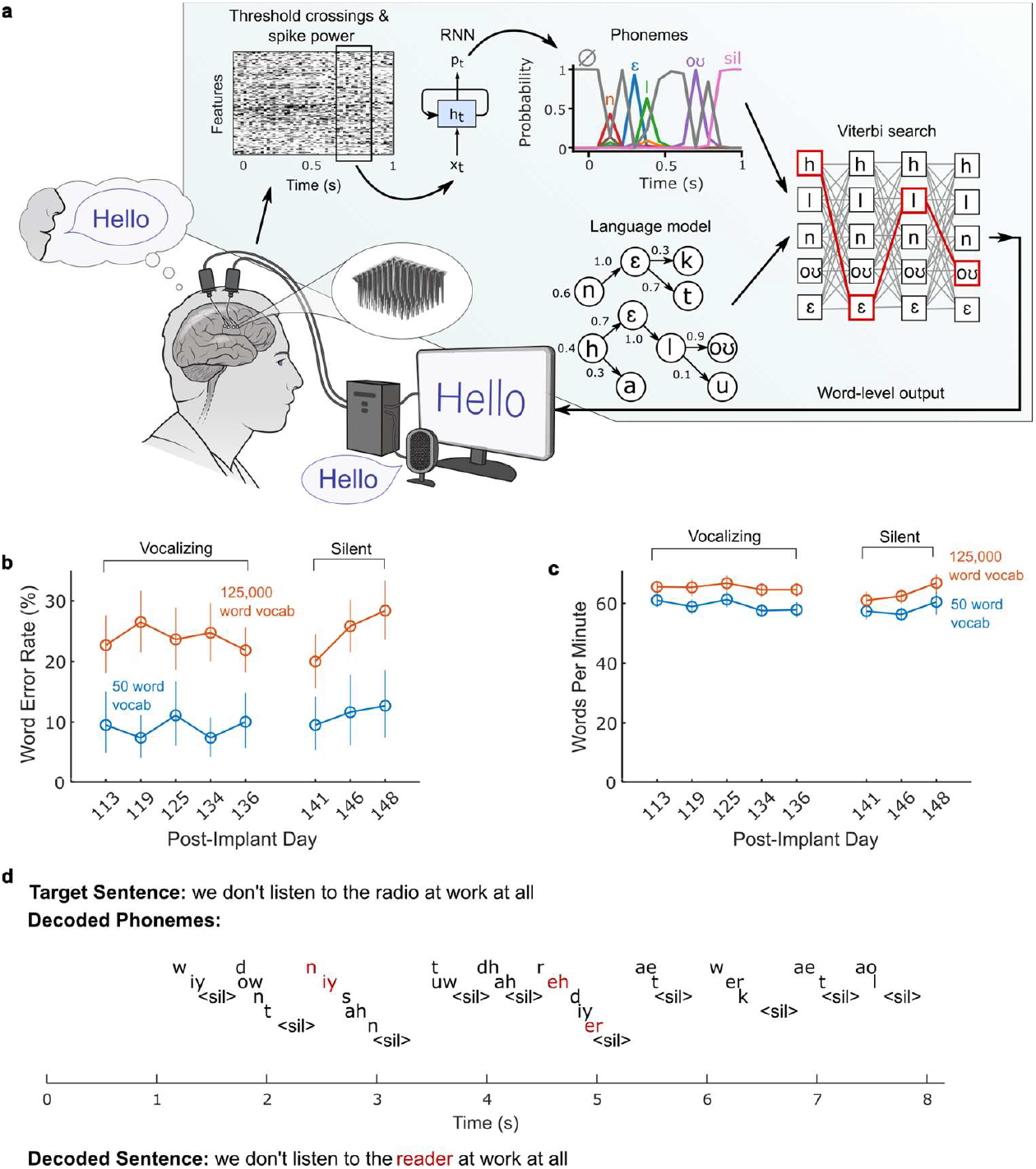
**(a)** Diagram of the decoding algorithm. First, the neural activity (multiunit threshold crossings and spike band power) is temporally binned and smoothed on each electrode. Then, a recurrent neural network (RNN) converts a time series of this neural activity into a probability for each phoneme (plus the probability of an interword “silence” token and a “blank” token associated with the connectionist temporal classification training procedure). The RNN is a 5-layer gated recurrent unit architecture trained using TensorFlow 2. Finally, the phoneme probabilities are combined with a large-vocabulary language model (a custom, 125,000-word trigram model implemented in Kaldi) to decode the most likely sentence. **(b)** Word error rates (edit distances) are shown for two speaking modes (vocalized vs. silent) and vocabulary sizes (50 vs 125,000 words). Vertical lines indicate 95% CIs. **(c)** Same as in b, but for speaking rate (words per minute). **(d)** A closed-loop example trial is shown, demonstrating the RNN’s ability to decode sensible sequences of phonemes (represented in ARPABET notation) without a language model. Phonemes are offset vertically for readability and “<sil>“ indicates the “silence” token (which the RNN was trained to produce at the end of all words). The phoneme sequence was generated by taking the maximum probability phonemes at each time step. Note that phoneme decoding errors are often corrected by the language model, which still infers the correct word. Incorrectly decoded phonemes and words are denoted in red.

At the beginning of each RNN performance evaluation day, we first recorded training data where T12 attempted to speak 260– 480 sentences at her own pace (41 ± 3.7 minutes of data; sentences were chosen randomly from the switchboard corpus^20^ of spoken English). A computer monitor cued T12 when to begin speaking and what sentence to speak. The RNN was then trained on this data in combination with all prior days’ data, using custom machine learning methods adapted from modern speech recognition^21–23^ to achieve high performance on limited amounts of neural data. In particular, we used unique input layers for each day to account for across-day changes in the neural activity, and rolling feature adaptation to account for within-day changes (SFig 5 highlights the effect of these and other architecture choices). By the last day, our training dataset consisted of 10,850 total sentences. Data collection and RNN training lasted 140 minutes per day on average (including breaks).

After training, the RNN was evaluated in real-time on held-out sentences that were never duplicated in the training set. For each sentence, T12 first prepared to speak the sentence during an instructed delay period. When the go cue was given, neural decoding was automatically triggered to begin. As T12 attempted to speak, neurally decoded words appeared on the screen in real-time reflecting the language model’s current best guess (SVideo 1). When T12 was finished speaking, she pressed a button to finalize the decoded output. We used two different language models: a large vocabulary model with 125,000 words (suitable for general English) and a small vocabulary model with 50 words (suitable for expressing some simple sentences useful in daily life). Sentences from the switchboard corpus^20^ were used to evaluate the RNN with the 125,000 word vocabulary. For the 50 word vocabulary, we used the word set and test sentences from Moses et al. 2021^2^.

Performance was evaluated over 5 days of attempted speaking with vocalization and 3 days of attempted silent speech (“mouthing” the words with no vocalization, which T12 reported she preferred because it was less tiring). Performance was consistently high for both speaking modes (Fig 2b-c, Table 1). T12 achieved a 9.1% word error rate for the 50 word vocabulary across all vocalizing days (11.2% for silent), and a 23.8% word error rate for the 125k word vocabulary across all vocalizing days (24.7% for silent). To our knowledge, this is the first successful demonstration of large-vocabulary decoding, and is also a significant advance in accuracy for small vocabularies (2.7 times fewer errors than prior work^2^). These accuracies were achieved at high speeds: T12 spoke at an average pace of 62 words per minute, which more than triples the speed of the prior state of the art for any type of BCI (18 words per minute for a handwriting BCI^8^).

**Table 1.** Mean phoneme and word error rates (with 95% CIs) for the speech BCI across all evaluation days. Phoneme error rates quantify the quality of the RNN decoder’s output before a language model is applied, while word error rates quantify the quality of the combined RNN and language model pipeline. Confidence intervals (CIs) were computed with the bootstrap percentile method (resampling over trials 10,000 times). “Online” refers to what was decoded in real-time, while “offline” refers to a post-hoc analysis of the data using an improved language model (“Improved LM”) or different partitioning of training and testing data (“Proximal Test Set”). In the proximal test set, training sentences occur much closer in time to testing sentences, mitigating the effect of within-day neural nonstationarities.

|  | Phoneme Error Rate %<br>[95% CI] | Word Error Rate %<br>[95% CI] |
| --- | --- | --- |
| **Online** |  |  |
| 125k Vocal | 19.7 [18.6, 20.9] | 23.8 [21.8, 25.9] |
| 125k Silent | 20.9 [19.3, 22.6] | 24.7 [22.0, 27.4] |
| 50 Vocal | 21.4 [19.6, 23.2] | 9.1 [7.2, 11.2] |
| 50 Silent | 22.1 [19.9, 24.3] | 11.2 [8.3, 14.4] |
| **Offline** |  |  |
| 125k Improved LM | 19.7 [18.6, 20.9] | 17.4 [15.4, 19.5] |
| 125k Improved LM +<br>Proximal Test Set | 17.0 [15.7, 18.3] | 11.8 [9.8, 13.9] |

Encouragingly, the RNN often decoded sensible sequences of phonemes before a language model was applied (Fig 2c). Phoneme error rates computed on the raw RNN output were 19.7% for vocal speech (20.9% for silent; see Table 1) and phoneme decoding errors followed a pattern related to speech articulation, where phonemes that are articulated similarly were more likely to be confused by the RNN decoder (SFig 6). These results suggest that good decoding performance is not overly reliant on a language model.

We also examined how information about speech production was distributed across the electrode arrays (SFig 7). We found that, consistent with Fig. 1, the ventral 6v array appeared to contribute more to decoding. Nevertheless, both arrays were useful, and low word error rates could be achieved only by combining both (offline analyses showed a reduction in word error rate from 32% to 21% when adding the dorsal array to the ventral array).

Finally, we explored the ceiling of decoding performance offline by (1) making further improvements to the language model and (2) evaluating the decoder on test sentences that occur closer in time to the training sentences (to mitigate the effects of within-day changes in the neural features across time). We found that an improved language model could decrease word error rates from 23.8% to 17.4%, and that testing on more proximal sentences further decreased word error rates to 11.8% (Table 1). These results indicate that substantial gains in performance are likely still possible with further language model improvements and more robust decoding algorithms that generalize better to nonstationary data (for example, unsupervised methods that track non-stationarities without requiring new training data ^24–27^).

### Preserved representation of speech articulation

Next, we investigated the representation of phonemes in area 6v during attempted speech. This is a challenging problem since we do not have ground truth knowledge of when each phoneme is being spoken (since T12 cannot speak intelligibly). To estimate how each phoneme was neurally represented, we analyzed our RNN decoders to extract vectors of neural activity (“saliency” vectors) that maximized the RNN probability output for each phoneme. We then asked whether these saliency vectors encode details about how phonemes are articulated.

First, we compared the neural representation of consonants to their articulatory representation, as measured by electromagnetic articulography in able-bodied speakers. We found broadly similar structure, which is especially apparent when ordering consonants by place of articulation (Fig 3A); the correlation between EMA and neural data was 0.61, far above chance (p<1e-4, Fig 3B). More detailed structure can also be seen – for example, nasal consonants are correlated (M, N, NG), and W is correlated with both labial consonants and velar/palatal consonants (since it contains aspects of both). Examining a low-dimensional representation of the geometry of the neural representation and articulatory representation shows a close match in the top two dimensions (Fig. 3c).

**Figure 3.**
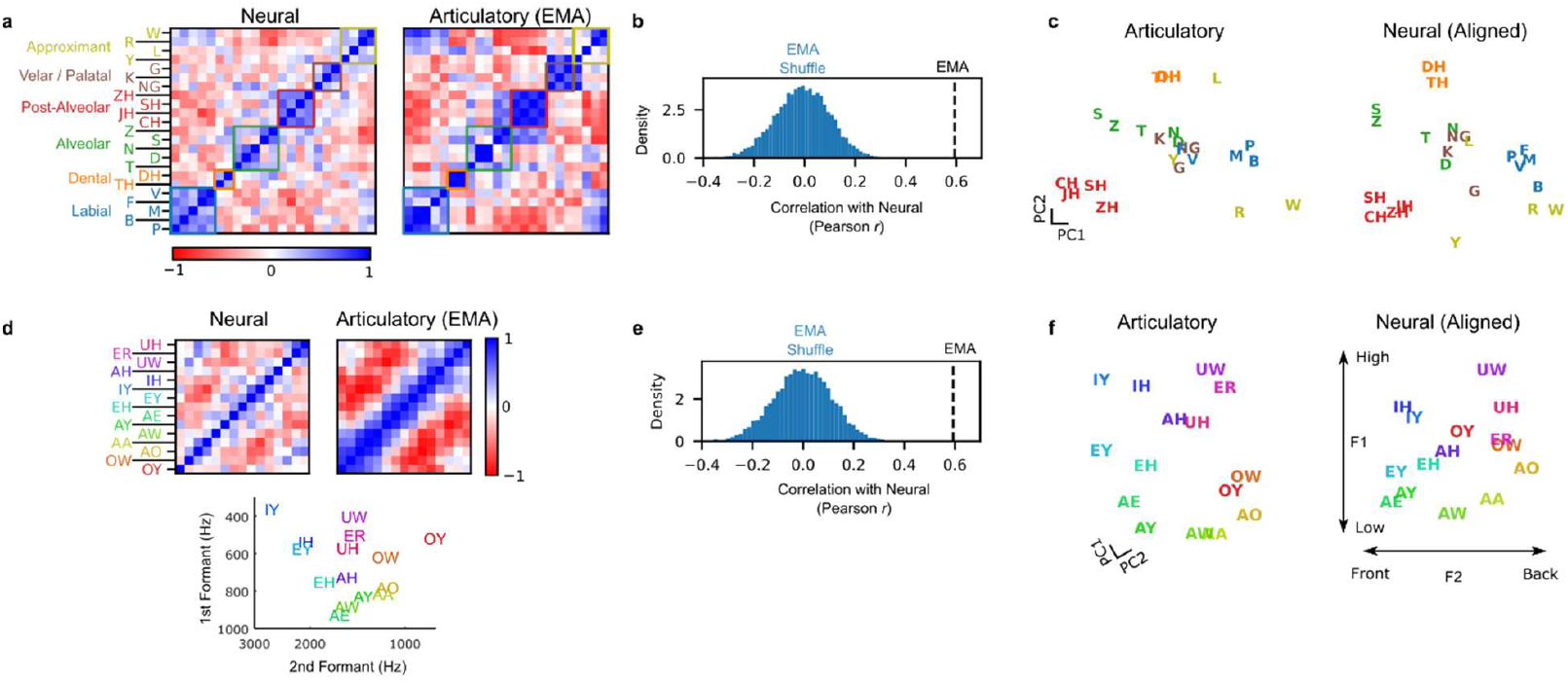
Preserved articulatory representation of phonemes. (a) Representational similarity across consonants for neural data (left) and articulatory data from an example subject who can speak normally obtained from the USC-TIMIT database (right). Each square in the matrix represents pairwise similarity for two consonants (as measured by cosine angle between the neural or articulatory vectors). Ordering consonants by place of articulation reveals a block-diagonal structure in the neural data that is also reflected in articulatory data. (b) Neural activity is significantly correlated with an articulatory representation. The blue distribution shows the correlations expected by chance. (c) Low-dimensional representation of phonemes articulatorily (left) and neurally (right). Neural data was rotated within the top 8 principal components (using cross-validated Procrustes) to show visual alignment with articulatory data. (d) Representational similarity for vowels, ordered by articulatory similarity. Diagonal banding in the neural similarity matrix indicates a similar neural representation. For reference, the first and second formant of each vowel is plotted below the similarity matrices^28^. (e) Neural activity correlates with the known two-dimensional structure of vowels. **(f)** Same as (c) but for vowels, with an additional within-plane rotation applied to align the (high vs. low) and (front vs. back) axes along the vertical and horizontal.

Next, we examined the representation of vowels, which have a two-dimensional articulatory structure: a high vs. low axis (how high the tongue is in the mouth, corresponding to the first formant frequency) and a front vs. back axis (whether the tongue is bunched up towards the front or back of the mouth, corresponding to the second formant frequency). We found that the saliency vectors for vowels mirror this structure, with vowels that are articulated similarly having a similar neural representation (Fig. 3d-e). Additionally, the neural activity contains a plane that reflects the two dimensions of vowels in a direct way (Fig. 3f).

Finally, we verified these results using additional ways of estimating neural and articulatory structure (SFig 8). Taken together, these results show that a detailed articulatory code for phonemes is still preserved even years after paralysis.

### Design considerations for speech BCIs

Finally, we examined three design considerations for improving the accuracy and usability of speech BCIs: language model vocabulary size, microelectrode count, and training dataset size.

To understand the effect of vocabulary size, we re-analyzed the 50-word-set data by re-processing the RNN output using language models of increasingly larger vocabulary sizes (Fig. 4a). We found that only very small vocabularies (e.g., 50– 100 words) retained the large improvement in accuracy relative to a large vocabulary model. Word error rates saturated at around 1,000 words, suggesting that using an intermediate vocabulary size may not be a viable strategy for increasing accuracy.

**Figure 4.**
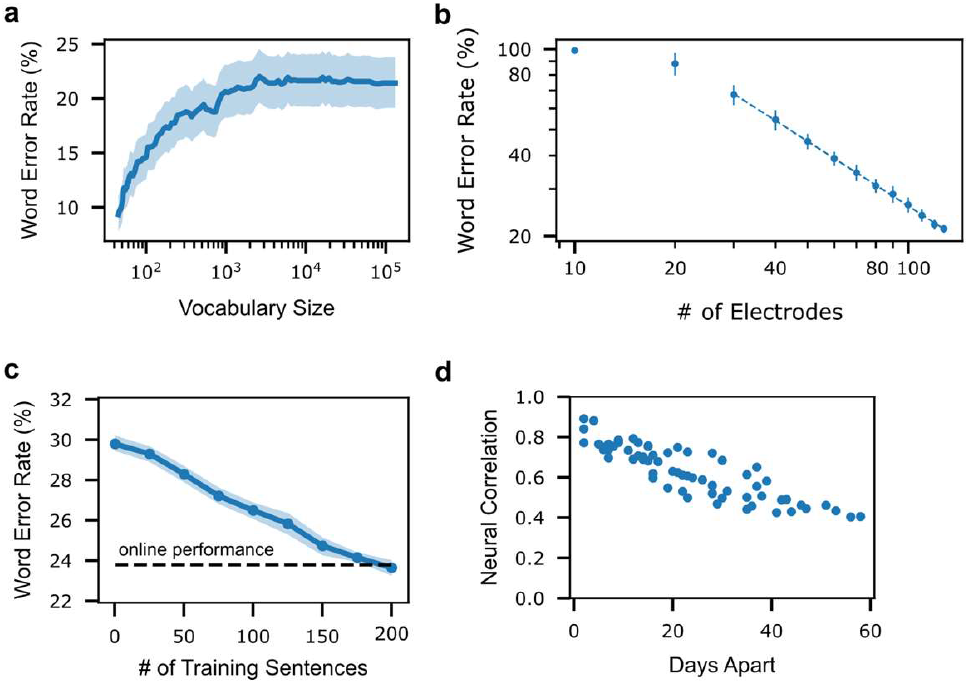
Design considerations for speech BCIs. (a) Word error rate as a function of language model vocabulary size, obtained by re-processing the 50-word set RNN outputs with language models of increasingly large vocabulary sizes (shaded region = 95% confidence interval). (b) Word error rate as a function of number of electrodes included in an offline decoding analysis (each filled circle shows the average word error rate of RNNs trained with that number of electrodes, and each thin line shows the standard deviation across 10 RNNs). There appears to be a log-linear relationship between # of electrodes and performance, such that doubling the electrode count cuts word error rate by nearly half (factor of 0.57; dashed line shows the log-linear relationship fit with least squares).. (c) Evaluation data from the 5 vocalized speech evaluation days was re-processed offline using RNNs trained in the same way, but with fewer (or no) training sentences taken from the day on which performance is evaluated. Word error rates are reasonable even when no training sentences are used from the evaluation day (i.e., when training on prior days’ data only). The dashed line shows online performance for reference (23.8% WER). (d) The correlation (Pearson r) in neural activity patterns representing a diagnostic set of words is plotted for each pair of days, revealing high correlations for nearby days.

Next, we investigated how accuracy improved as a function of the number of electrodes used for RNN decoding. Accuracy improved monotonically with a log-linear trend (Fig. 4b – doubling the electrode account appears to cut the error rate nearly in half).This suggests that intracortical devices capable of recording from more electrodes (e.g., denser or more extensive microelectrode arrays) may be able to achieve improved accuracies in the future, although the extent to which this downward trend will continue remains to be seen..

Finally, in this demonstration we used a large amount of training data per day (260 – 440 sentences). Retraining the decoder each day helps to adapt to neural changes that occur across days. We examined offline if this much data per day was necessary, by re-processing the data with RNNs trained in the same way but with fewer sentences. We found that performance was good even without using any training data on the new day (Fig. 4c, word error rate = 30% with no retraining). Furthermore, we found that neural activity changed at a gradual rate over time, suggesting that unsupervised algorithms for updating decoders to neural changes should be feasible (Fig. 4d) ^24–27^.

## Discussion

People with neurological disorders such as brainstem stroke or amyotrophic lateral sclerosis (ALS) frequently face severe speech and motor impairment and, in some cases, completely lose the ability to speak (locked-in syndrome^29^). Recently, BCIs based on hand movement activity have enabled typing speeds of 8-18 words per minute in people with paralysis^8,30^. Speech BCIs have the potential to restore natural communication at a much faster rate, but have not yet achieved high accuracies on large vocabularies (i.e., unconstrained communication of any sentence the user may want to say)^1–7^. Here, we demonstrated a speech BCI that can decode unconstrained sentences from a large vocabulary at a speed of 62 words per minute, using microelectrode arrays to record neural activity at single neuron resolution. To our knowledge, this is the first time that a BCI has substantially exceeded the communication rates that alternative technologies can provide for people with paralysis (e.g., eye tracking^31^).

Our demonstration is a proof of concept that decoding attempted speaking movements with a large vocabulary is possible using neural spiking activity. However, it is important to note that it does not yet constitute a complete, clinically viable system. Work remains to be done to reduce the time needed to train the decoder and adapt to changes in neural activity that occur across days without requiring the user to pause and recalibrate the BCI (see ^32,24–27^ for initial promising approaches). Additionally, intracortical microelectrode array technology is still maturing ^33,34^, and is expected to require further demonstrations of longevity and efficacy before widespread clinical adoption (although recent safety data is encouraging^35^, and next-generation recording devices are under development^36,37^). Furthermore, the decoding results shown here must be confirmed in additional participants, and their generalizability to people with more profound orofacial weakness remains an open question. Variability in brain anatomy is also a potential concern, and more work must be done to confirm that regions of precentral gyrus containing speech information can be reliably targeted.

Perhaps most importantly, a 24% word error rate is likely not yet low enough for everyday use (e.g., compare to 4-5% word error rate for state of the art speech-to-text systems^23,38^). Nevertheless, we believe that our results show promise for decreasing the word error rates further. First, word error rate decreases as more channels are added, suggesting that intracortical technologies that record more channels may enable lower word error rates in the future. Second, room still remains for optimizing the decoding algorithm; with further language model improvements and when mitigating the effect of within-day nonstationarities, we were able to reduce the word error rate to 11.8% in offline analyses. Finally, we showed that ventral premotor cortex (area 6v) contains a rich, intermixed representation of speech articulators even within a small area (3.2 × 3.2 mm), and that the details of how phonemes are articulated are still faithfully represented even years after paralysis in someone who can no longer speak intelligibly. Taken together, this suggests that a higher channel count system that records from only a small area of 6v is a feasible path forward towards a device that can restore communication at conversational speeds to people with paralysis.

## Acknowledgments

We thank participant T12 and her caregivers for their generously volunteered time and effort as part of the BrainGate2 pilot clinical trial, Beverly Davis, Kathy Tsou, and Sandrin Kosasih for administrative support, Yaguang Hu for providing suggestions about the language model, and Matt Glasser, Tim Coalson, David Van Essen and B. Choi for help with the HCP pipeline. Support provided by the Office of Research and Development, Rehabilitation R&D Service, Department of Veterans Affairs (N2864C, A2295R); Wu Tsai Neurosciences Institute; Howard Hughes Medical Institute; Larry and Pamela Garlick; Simons Foundation Collaboration on the Global Brain; NIDCD U01-DC017844, NIDCD U01-DC019430 (algorithm development only).

---

**Supplemental Video 1**. In this video, participant T12 uses the speech BCI in real-time to copy sentences shown on the screen. When the square in the center of the screen is red, T12 reads the sentence above the square and prepares to speak it. When the square turns green, T12 attempts to speak that sentence while the real-time decoder output is shown below the square. Note that T12 produces unintelligible vocalizations when attempting to speak. When T12 is finished speaking, a text-to-speech program reads the final decoded text aloud. These sentences were recorded during a performance evaluation session reported in Figure 2 (post-implant day 136).

**Supplemental Video 2**. The same as supplemental video 1, except T12 is silently speaking (i.e., mouthing the words) instead of attempting to produce vocalizations. These sentences were recorded on post-implant day 141.

**Supplemental Figure 1.**
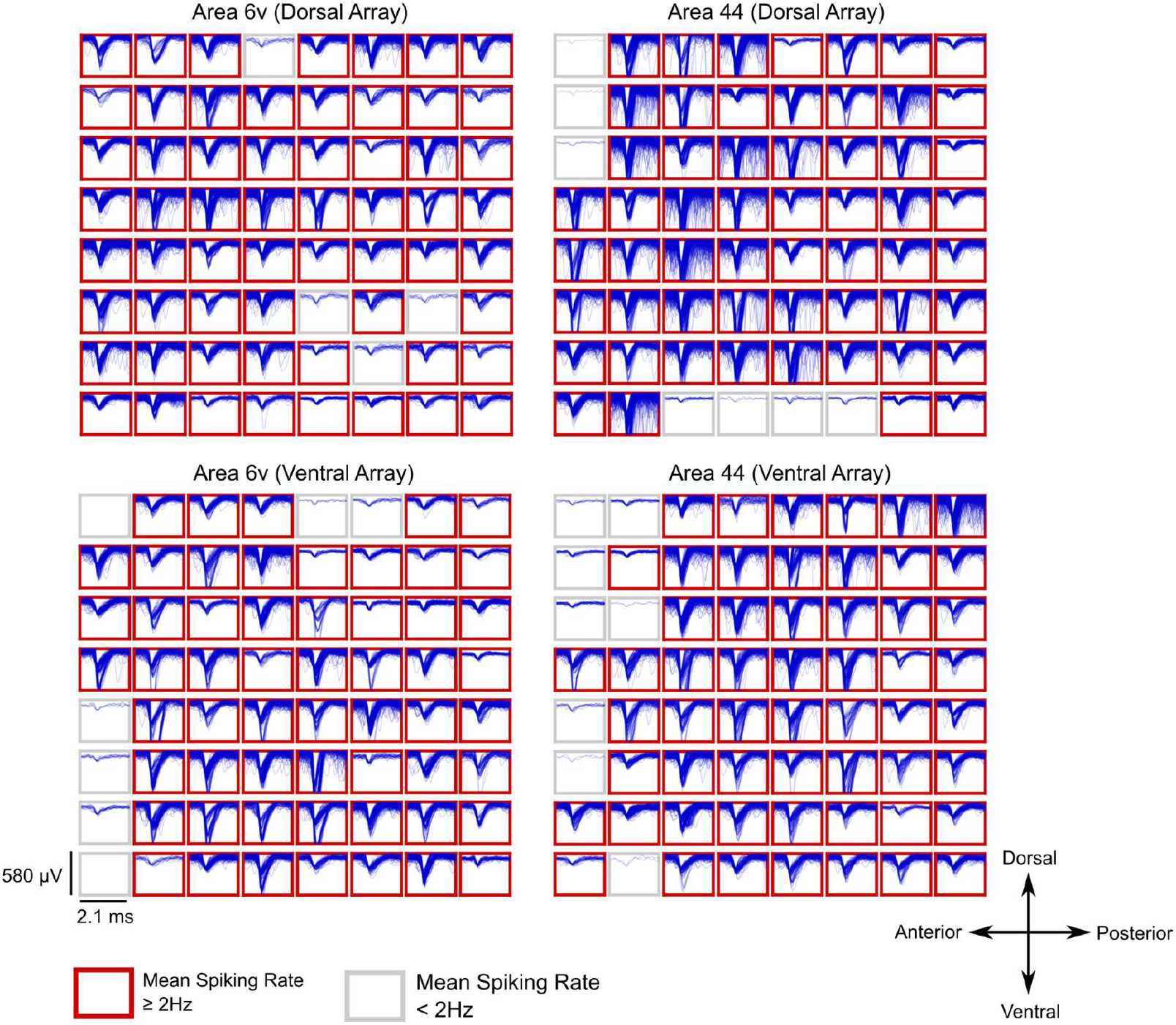
Example spike waveforms detected during a 10-s time window are plotted for each electrode (data were recorded on post-implant day 119). Each 8×8 grid corresponds to a single 64-electrode array, and each rectangular panel in the grid corresponds to a single electrode. Blue traces show example spike waveforms (2.1-ms duration). Neural activity was band-pass filtered (250-5000 Hz) with an acausal, 4^th^ order Butterworth filter. Spiking events were detected using a −4.5 root mean square (RMS) threshold, thereby excluding almost all background activity. Electrodes with a mean threshold crossing rate of at least 2 Hz were considered to have ‘spiking activity’ and are outlined in red (note that this is a conservative estimate that is meant to include only spiking activity that could be from single neurons, as opposed to multiunit ‘hash’). The results show that many electrodes record large spiking waveforms that are well above the noise floor (the *y* axis of each panel spans 580 μV, whereas the background activity has an average RMS value of only 30.8 μV). In area 6v,118 electrodes out of 128 had a threshold crossing rate of at least 2 Hz on this particular day (113 electrodes out of 128 in area 44).

**Supplemental Figure 2.**
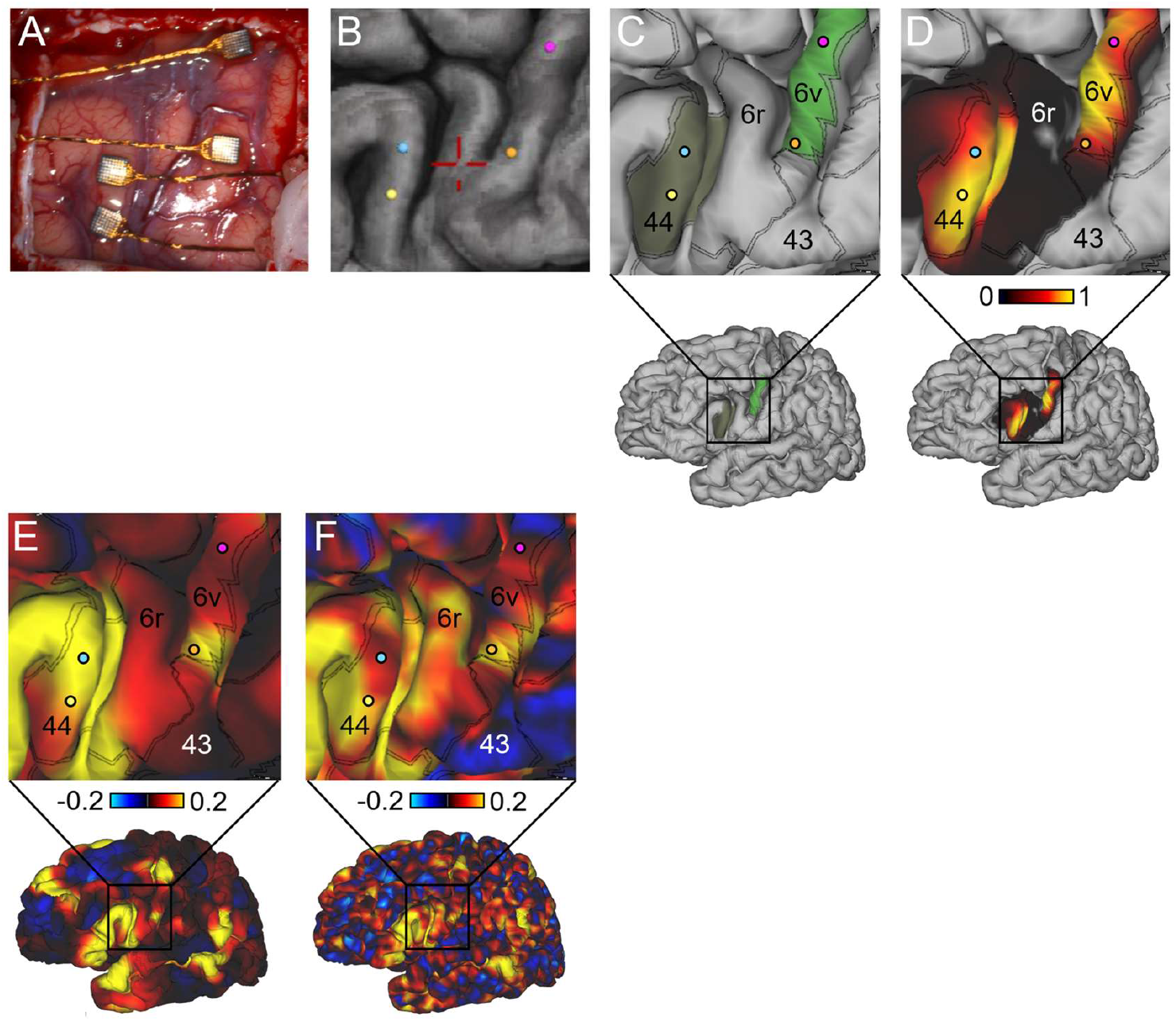
Locations of the implanted arrays within areas 6v and 44 are shown (A) directly on the participant’s brain surface and (B) on a 3D reconstruction of the brain and the implanted arrays (array centers shown in blue, yellow, magenta, and orange circles) in StealthStation (Medtronic, Inc.). (C) shows the approximate array locations in the participant’s inflated brain using Connectome Workbench software, overlaid on the cortical areal boundaries (double black lines) estimated by the Human Connectome Project (HCP) cortical parcellation. (D) shows the approximate array locations on the same brain as (C) overlaid on the confidence maps of the areal regions. (E,F) A language-related resting state network identified in the Human Connectome Project data (N=210) aligned to T12’s brain (E) and in T12’s individual scan (F). The ventral part of 6v appears to be involved in this resting state network, while the dorsal part is not. This resting state network includes language-related area 55b, Broca’s area and Wernicke’s area.

**Supplemental Figure 3.**
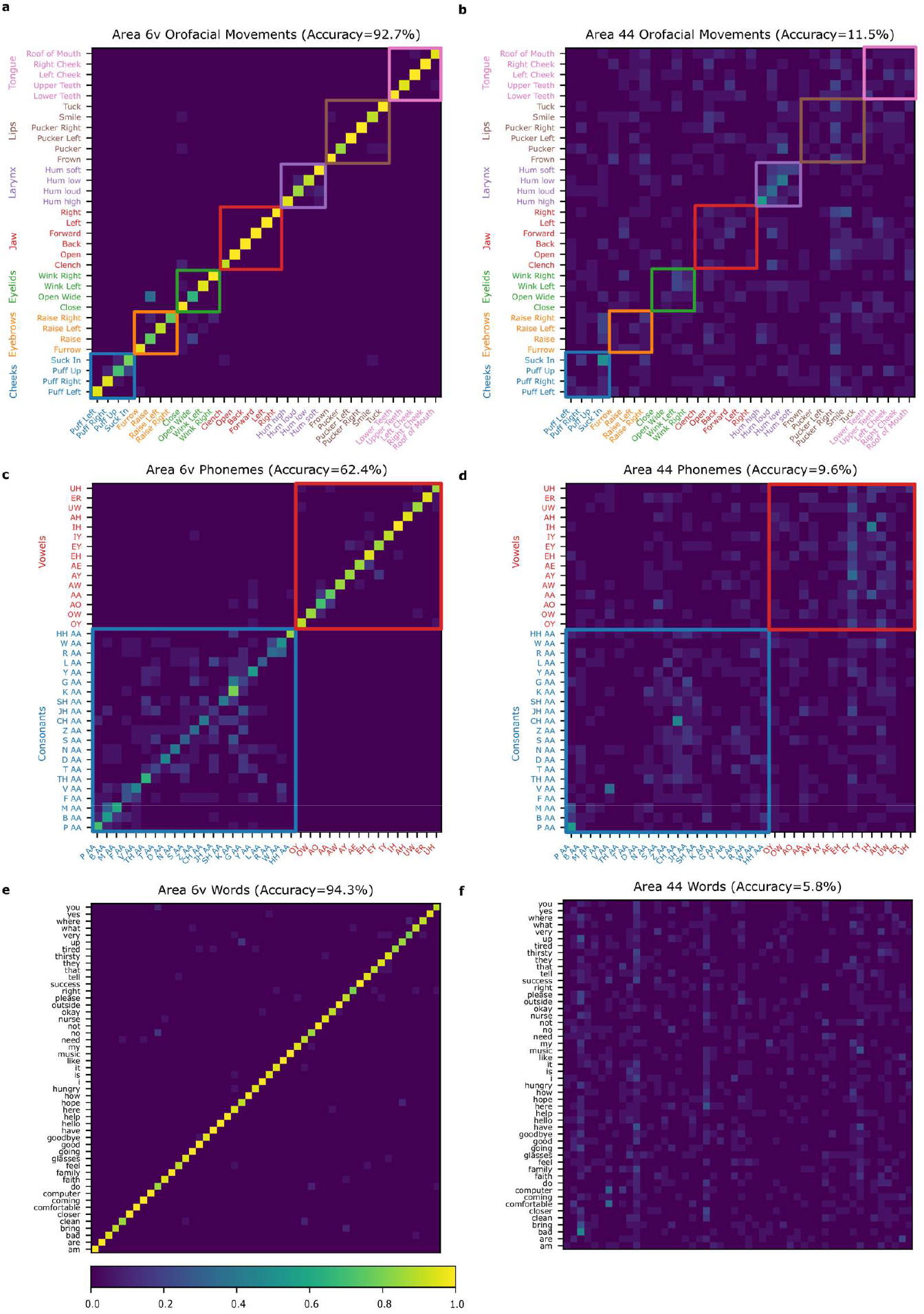
Confusion matrices from a cross-validated, Gaussian naïve Bayes classifier trained to classify amongst orofacial movements (a-b), individual phonemes (c-d), or individual words (e-f) using threshold crossing rates averaged in a window from 0 to 1000 ms after the go cue. Each entry (i, j) in the matrix is colored according to the percentage of trials where movement j was decoded (of all trials where movement i was cued). Matrices (a,c,e) show results from using only electrodes in area 6v, while matrices (b,d,f) show results from using electrodes in area 44. While good classification performance can be observed from area 6v, area 44 appears to contain little to no information about most movements, phonemes and words.

**Supplemental Figure 4.**
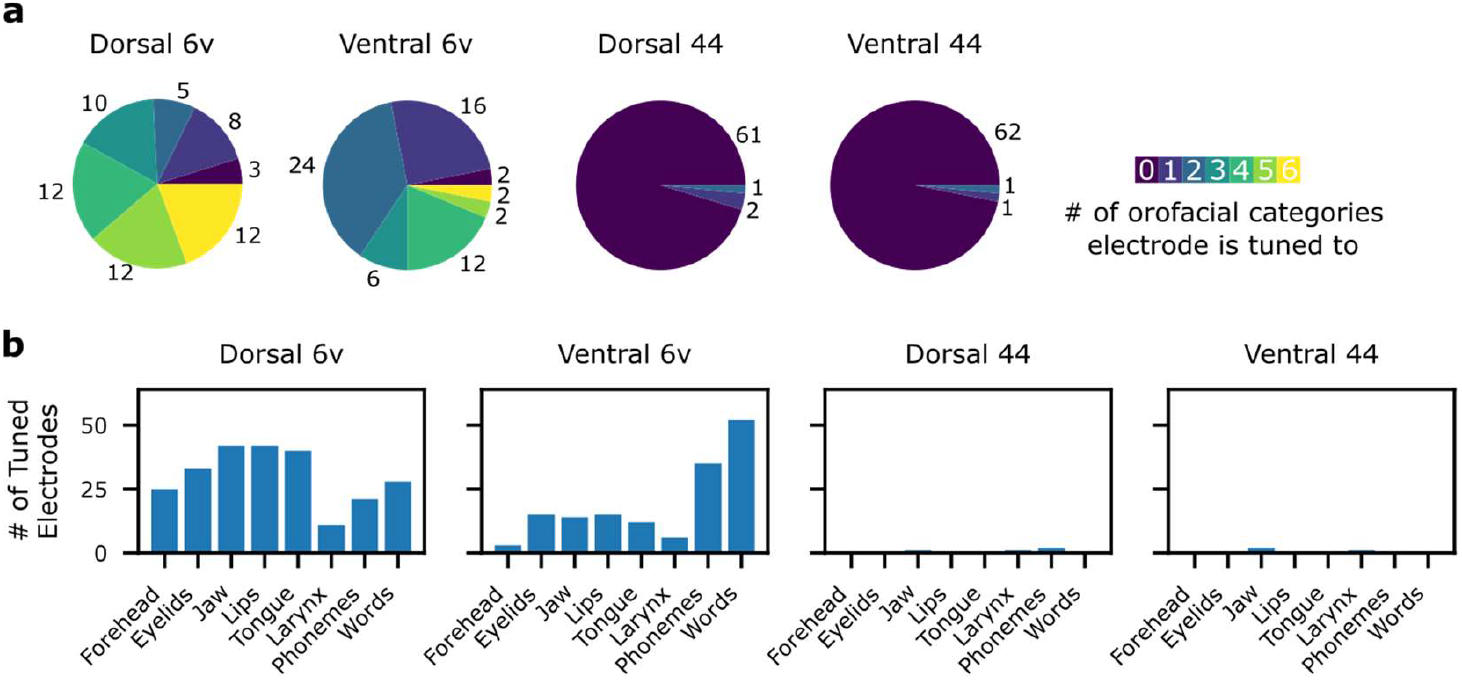
(a) Pie charts summarizing the number of electrodes that had statistically significant tuning to each possible number of movement categories (from 0 to 6), as assessed with a 1-way ANOVA (p<1e-5). On the 6v arrays, many electrodes are tuned to more than one orofacial movement category (forehead, eyelids, jaw, lips, tongue, and larynx). (b) Bar plots summarizing the number of tuned electrodes to each movement category and each array. The ventral 6v array contains more electrodes tuned to phonemes and words, while the dorsal 6v array contains more electrodes tuned to orofacial movement categories. Nevertheless, both 6v arrays contain electrodes tuned to all categories of movement.

**Supplemental Figure 5.**
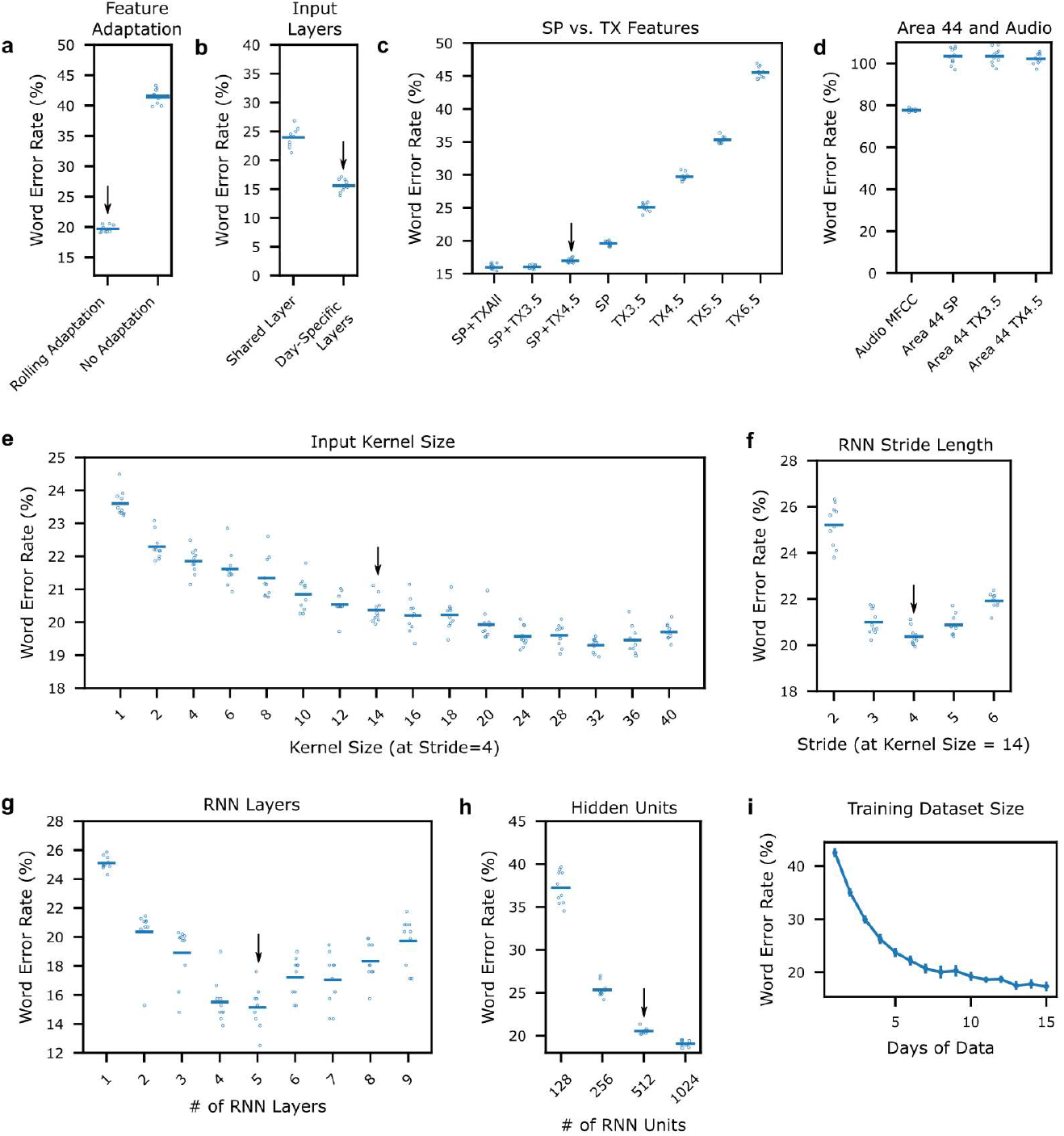
Effect of RNN parameters and architecture choices, as determined by evaluating RNN performance in offline parameter sweeps. Black arrows denote the parameters used for real-time evaluation. Each blue open circle denotes the performance of a single RNN seed, and each thin blue bar denotes the mean across all seeds from that parameter configuration. (a) Rolling z-scoring improves performance substantially as compared to no feature adaptation (when testing on held-out blocks that are separated in time from the training data). (b) Training RNNs with unique (day-specific) input layers improves performance relative to using a shared layer that is the` same across all days. (c) RNN performance using different neural features as input (SP=spike band power, TX=threshold crossing). Combining spike band power with threshold crossings performs better than either one alone, and it appears that performance could have been improved slightly by using a -3.5 RMS threshold instead of - 4.5. (d) RNN performance using audio mel frequency cepstral coefficients (audio MFCC) or neural features from the area 44 arrays (IFG). While poor but above-chance performance can be achieved using MFCCs, word error rates from IFG recordings appear to be at chance level (∽100%). (e) RNN performance as a function of “kernel size” (i.e., the number of 20 ms bins stacked together as input and fed into the RNN at each time step). It appears that performance could have been improved by using larger kernel sizes. (f) RNN performance as a function of “stride” (a stride of *N* means the RNN steps forward only every *N* time bins). (g) RNN performance as a function of the number of stacked RNN layers. (h) RNN performance as a function of the number of RNN units per layer. (i) RNN performance as a function of the number of prior days included as training data. Performance improves by adding prior days, but with diminishing returns. The blue line shows the average word error rate across 10 RNN seeds and 5 evaluation days. Vertical lines show standard deviations across the 10 seeds.

**Supplemental Figure 6.**
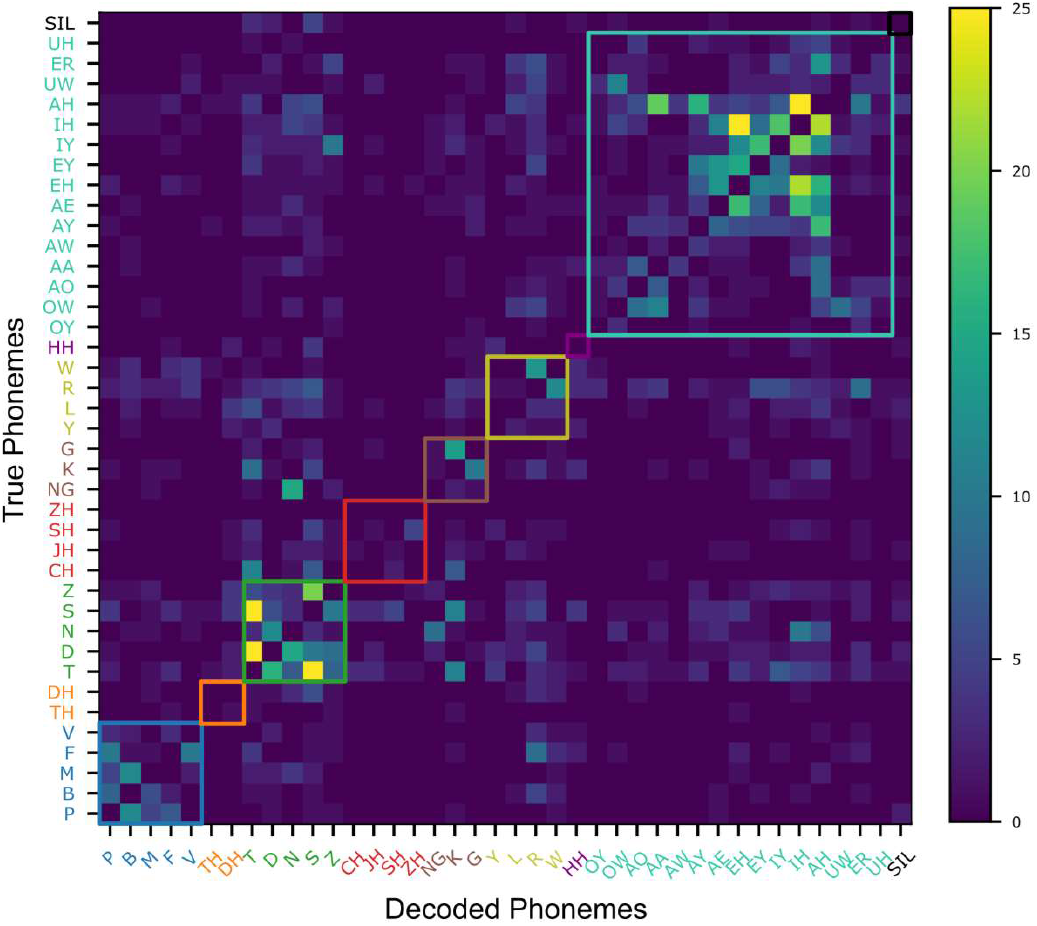
Phoneme substitution errors observed across all real-time evaluation sentences. Entry *(i.j)* in the matrix represents the substitution count observed for true phoneme *i* and decoded phoneme *j*. Substitutions were identified using an edit distance algorithm that determines the minimum number of insertions, deletions, and substitutions required to make the decoded phoneme sequence match the true phoneme sequence. Most substitutions appear to occur between phonemes that are articulated similarly.

**Supplemental Figure 7.**
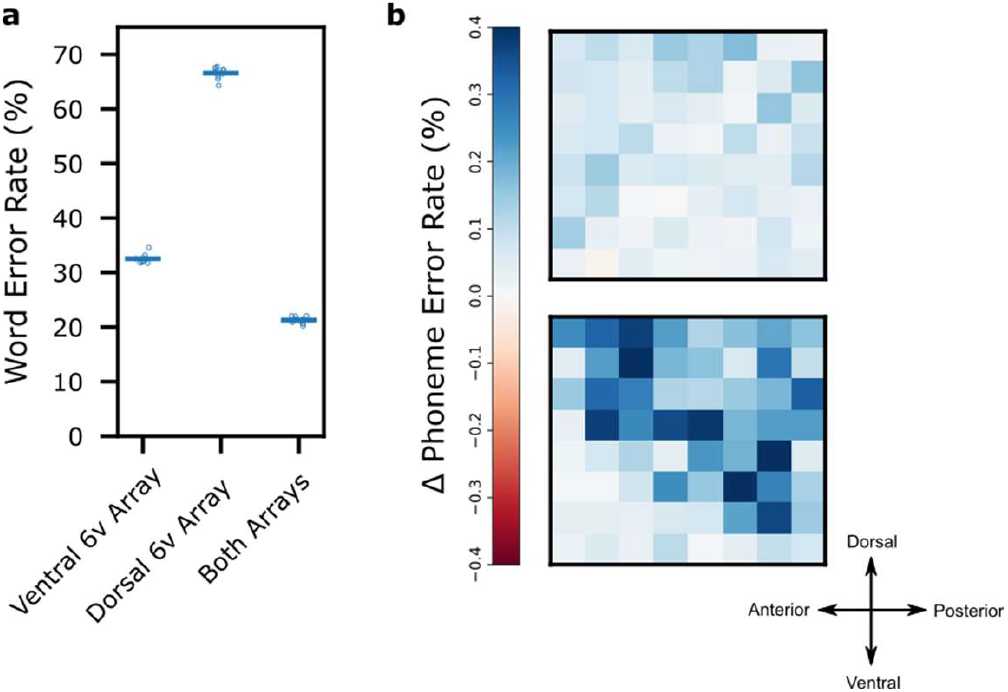
(a) Word error rates of a retrospective offline decoding analysis using just the ventral 6v array (left column), dorsal 6v array (middle column), or both arrays (right column). (b) Heatmaps depicting the (offline) increase in phoneme error rate when removing each electrode from the decoder by setting the values of its corresponding features to zero (averaged across 10 RNN seeds that were all originally trained to use every electrode). Almost all electrodes seem to contribute to decoding performance, although the most informative electrodes are concentrated on the ventral array. The effect of removing any one electrode is small (<1% increase in phoneme error rate), owing to the redundancy across electrodes.

**Supplemental Figure 8.**
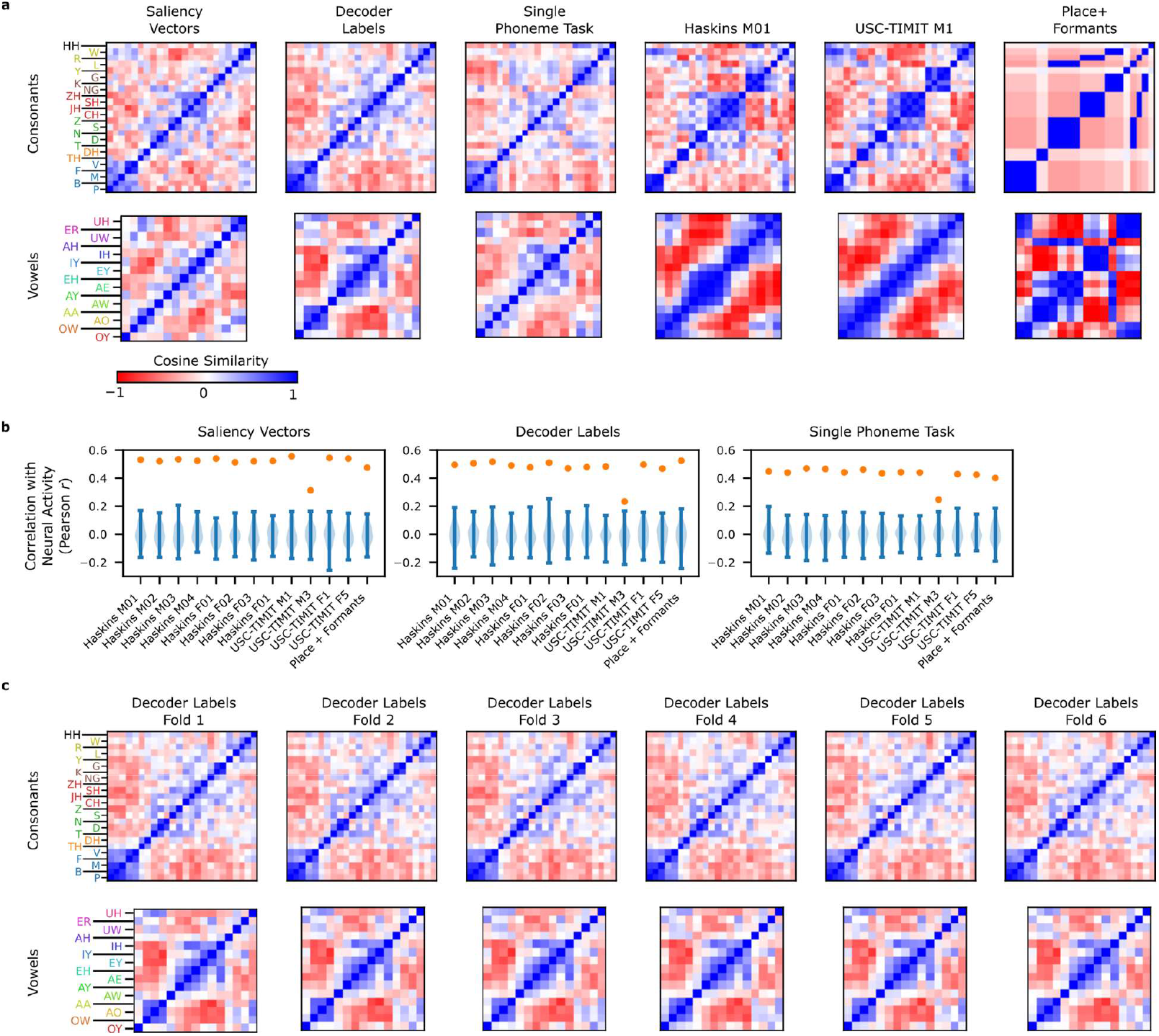
(a) Different methods of quantifying articulatory structure in neural activity and articulator kinematics all yield similar results. (a) Representational similarity across consonants (top) and vowels (bottom) for different quantifications of the neural activity (“Saliency Vectors”, “Decoder Labels”, and “Single Phoneme Task”) and articulator kinematics (“Haskins M01”, “USC-Timit M1”, “Place + Formants”). Each square in a matrix represents pairwise similarity for two phonemes (as measured by cosine angle between the neural or articulatory vectors). Consonants are ordered by place of articulation (but with approximants grouped separately) and vowels are ordered by articulatory similarity (as measured by “USC-TIMIT M1”). These orderings reveal block-diagonal structure in the neural data that is also reflected in articulatory data. “Haskins M01” and “USC-Timit M1” refer to subjects M01 and M1 in the Haskins and USC-Timit datasets. “Place + Formants” refers to coding consonants by place of articulation and to representing vowels using their two formant frequencies. (b) Correlations between the neural representations and the articulator representations (each panel corresponds to one method of computing the neural representation, while each column corresponds to one EMA subject or the place/formants method). Orange dots show the correlation value (Pearson *r*), and blue distributions show the null distribution computed with a shuffle control (10,000 repetitions). In all cases, the true correlation lies outside the null distribution (p<1e-4). Correlation values were computed between consonants and vowels separately and then averaged together to produce a single value. (c) Representational similarity matrices computed using the “Decoder labels” method on 6 different independent folds of the neural data. Very similar representations across folds indicates that the representations are statistically robust (average correlation across folds = 0.79).

## 1. Experimental procedures

### 1.1. Study participant

This study includes data from one participant (identified as T12) who gave informed consent and was enrolled in the BrainGate2 Neural Interface System clinical trial (ClinicalTrials.gov Identifier: NCT00912041, registered June 3, 2009). This pilot clinical trial was approved under an Investigational Device Exemption (IDE) by the US Food and Drug Administration (Investigational Device Exemption #G090003). Permission was also granted by the Institutional Review Board of Stanford University (protocol #52060). T12 gave consent to publish photographs and videos containing her likeness. All research was performed in accordance with relevant guidelines/ regulations.

T12 is a left-handed woman, 67 years old at the time of data collection, with slowly-progressive bulbar-onset Amyotrophic Lateral Sclerosis (ALS) diagnosed at age 59 (ALS-FRS score of 26 at the time of study enrollment). On March 30, 2022, four 64-channel, 1.5 mm-length silicon micro electrode arrays coated with sputtered iridium oxide (Blackrock Microsystems, Salt lake City, UT) were implanted in T12’s left hemisphere, based on preoperative anatomical and functional magnetic resonance imaging (MRI) and cortical parcellation (see sections 1.2 and 1.3 below for details). Two arrays were placed in area 6v (oral-facial motor cortex) of ventral precentral gyrus, and two were placed in area 44 of inferior frontal gyrus (considered part of Broca’s area). Data are reported from post-implant days 27-148. On average, 119.6 ± 5.0 (Mn ± sd) out of 128 electrodes recorded spike waveforms at a rate of at least 2 Hz when using a spike-detection threshold of -4.5 RMS, where RMS is the electrode-specific root mean square of the voltage time series recored on that electrode (see SFig 1 for example waveforms).

T12 is severely dysarthric due to bulbar ALS and has been for nearly 8 years. She retains partial use of her limbs, and communicates primarily through use of a writing board or iPad tablet. She is able to vocalize while attempting to speak, and is able to produce some subjectively differentiable vowels sounds. However, we had difficulty discerning nearly all consonants produced in isolation (with the possible exception of the bilabial nasal consonant “M”), and could not reliably make out any consonants or vowels when T12 attempted to speak whole sentences at a fluent rate (SVideo 1 shows examples of attempted speaking).

### 1.2. Functional MRI speech lateralization

Prior to surgery, participant T12 underwent anatomic and functional brain imaging on a GE Discovery MR750 3T MRI scanner, using a routine clinical acquisition protocol, in order to determine whether she was right or left hemisphere dominant for language. BOLD fMRI images were acquired using T2*-weighted volumes collected with 4 mm slice thickness and 2 ×2 mm^2^ in-plane voxel resolution. BOLD images were acquired during performance of a suite of tasks including visually responsive naming, object naming, auditory responsive naming, repetitive movements of the right hand, left hand, right foot, left foot, and tongue. Tasks were performed in 4 minute blocks consisting of repeated sequences of 10 seconds of task performance followed by 10 seconds of rest. Task instructions were presented by the SensaVue presentation system with verbal instructions given by the MRI technologist. T-score thresholds for processing were chosen based on direct inspection of the preliminary fMRI output for each task produced by the scanner software and inspected by the interpreting radiologist at the scanner.

Following scan completion, the complete data set was sent to and processed by DynaSuite. Fully processed fMRI statistical parametric maps for each task were registered to and overlaid on an anatomic 3D gradient echo T1-weighted (BRAVO) image acquired with 1 mm isotropic voxels. Quality control steps including motion tracking and assessment of anatomic-functional registration fidelity. Language lateralization was assessed by visual inspection of lateralization of activation in Broca’s area, Wernicke’s area, speech supplemental motor area, and the basal temporal language area. Results indicated a clear left hemisphere lateralization of language in T12.

### 1.3. Array placement targeting

The surgical targets for array placement within areas 6v and 44 were selected based on gross anatomical structure (e.g., gyri and sulci), vasculature, and estimates of the boundaries of areas 44 and 6v obtained using a cortical parcellation method derived from multi-modal Human Connectome Project (HCP) data [1].

We acquired T1-weighted (T1w), T2-weighted (T2w), resting-state functional MRI (rsfMRI), single band fMRI, and spin echo fieldmap images to generate the HCP-based cortical parcellation. The participant was scanned in a 3T Ultra High Performance scanner (GE Healthcare) with a Nova 32-channel coil. Scan parameters were based on HCP Lifespan protocols and modified for the GE system (Table 1).

Data were processed using the HCP pipelines as described on https://github.com/Washington-University/HCPpipelines (see Glasser et al. 2016 for further details). Briefly, T1w and T2w images were initially preprocessed using the FreeSurfer pipeline (version 7.1.1) to perform motion, distortion, and bias field corrections; brain extraction; white matter segmentation; cortical surface reconstruction, and spherical mapping (PreFreeSurferPipelineBatch.sh, FreeSurferPipelineBatch.sh). Surface outputs were then aligned to the standard surface template using MSMSulc, as well as used to create myelin maps (PostFreeSurferPipelineBatch.sh).

rsfMRI data were corrected for motion, bias field, and susceptibility distortions using the spin echo fieldmaps and single band reference fMRI images and non-linearly registered to MNI space (GenericfMRIVolumeProcessingPipelineBatch.sh), followed by volume to surface mapping (GenericfMRISurfaceProcessingPipelineBatch.sh). Data then underwent spatial MELODIC ICA (IcaFixProcessingBatch.sh), manual classification of the components as signal or noise, and denoising.

The MSMAll pipeline was then run to re-align the participant’s cortical surface to the standard surface template using areal features from the cortical folding map, myelin map, rsfMRI networks, and rsfMRI-based retinotopy. Lastly, the data also underwent a dedrift and resample step (DeDriftAndResamplePipelineBatch.sh). This then allowed us to overlay the Human Connectome Project (HCP) cortical parcellation (210P) and areal confidence (210V) maps [1] onto the participant’s brain using Connectome Workbench, thereby obtaining estimates of areas 6v and 44 within the participant’s brain (as shown in SFig 2). In SFig 2E-F we show resting state network 25 (an ICA component originally identified in [1] as the “Language RSN” shown there in supplemental figure 8a).

### 1.4. Neural signal processing

Neural signals were recorded from the microelectrode arrays using the Neuroplex-E system (Blackrock Microsystems) and transmitted via a cable attached to a percutaneous connector. Signals were analog filtered (4th order Butterworth with corners at 0.3 Hz to 7.5 kHz), digitized at 30 kHz (250 nV resolution), and fed to custom software written in Simulink (Mathworks) for digital filtering and feature extraction. Digital filtering began with a highpass filter (300 Hz cutoff) that was applied non-causally to each electrode, using a 4 ms delay, in order to improve spike detection [2]. Linear regression referencing (LRR) was then applied to further reduce reduce ambient noise artifacts [3].

After filtering, binned threshold crossing counts (20 ms bins) were computed by counting the number of times the filtered voltage time series crossed an amplitude threshold set at – 4.5 times the standard deviation of the voltage signal. Electrode-specific thresholds and LRR filter coefficients were set using data recorded from an initial “diagnostic” block at the beginning of each session (see section 1.7 for more details). Binned spike band power (20 ms bins) was computed by taking the sum of squared voltages observed during each time bin. Threshold crossing rates and spike band power are commonly used measurements of local spiking activity that have been shown to be comparable to sorted single unit activity in terms of decoding performance and neural population structure [4, 5, 6]. For decoding, threshold crossing counts and spike band power from the 128 electrodes in area 6v were concatenated to yield a 256 × 1 feature vector per time step. For neural tuning analyses (e.g. Figure 1), only threshold-crossing counts were used.

**Table 1.** MRI Scan Parameters for Cortical Parcellation.

| Image | T1w | T2w | rsfMRI | rsfMRI-single band | Spin echo fieldmap |
| --- | --- | --- | --- | --- | --- |
| Sequence | 3D MPRAGE | 3D CUBE | 2D Gradient Echo EPI | 2D Gradient Echo EPI | 2D Spin Echo EPI |
| TR (ms) | 3000 | 2500 | 800 | 4200 | 8000 |
| TE (ms) | 3.5 | 60-78 | 37 | 30 | min full |
| TI (ms) 1060 | - | - | - | - | - |
| Parallel imaging | 2 x 1.25 | 1.9 x 1.9 | - | - | - |
| Fat suppression | no | no | yes | yes | yes |
| Resolution (mm) | 0.8 x 0.8 x 0.8 | 0.8 x 0.8 x 0.8 | 2 x 2 x 2 | 2 x 2 x 2 | 2 x 2 x 2 |
| Matrix size | 320 x 320 x 230 | 320 x 320 x 216 | 104 x 104 x 72 | 104 x 104 x 72 | 104 x 104 x 72 |
| FOV (mm) | 256 x 256 x 184 | 256 x 256 x 184 | 208 x 208 x 144 | 208 x 208 x 144 | 208 x 208 x 144 |
| Flip angle | 8 | - | 54 | 90 | - |
| Slice orientation | sagittal, AC-PC | sagittal, AC-PC | axial AC-PC | axial AC-PC | axial AC-PC |
| Phase encoding | - | - | AP and PA (separately) | AP and PA (separately) | AP and PA (separately) |
| Multiband factor | - | - | 8 | 1 | - |

### 1.5. Data collection rig

Digital signal processing and feature extraction was performed on a dedicated computer using Simulink Real-Time. Extracted features were then sent to a separate computer running Ubuntu for neural decoding and recording. Decoding and recording software was written in Python using TensorFlow 2 and Redis. The Ubuntu computer also ran the experimental task software that displayed cues to T12 on a computer monitor. The task software was implemented using MATLAB and the Psychophysics Toolbox [7]). Finally, a third computer running Windows was used to interface with the Neuroplex-E system and control the starting and stopping of experimental tasks.

### 1.6. Overview of data collection sessions

Neural data were recorded in 2– 4 hour “sessions” on scheduled days, which typically occurred 2 times per week. During the sessions, T12 sat in either a wheelchair or power lift chair in an upright position, with a pillow placed to support her head and neck, and her hands resting on her lap. A computer monitor placed in front of T12 indicated which sentence to speak (or which movement to make) and when. Data were collected in a series of 5– 10 minute “blocks” consisting of an uninterrupted series of trials. In between these blocks, T12 was encouraged to rest as needed. Table 2 below lists all 26 data collection sessions reported in this work.

**Table 2:** Data Collection Sessions

| Session Number | Date (Post-Implant Day) | Description | Data |
| --- | --- | --- | --- |
| 1 | 2022.04.21 (27) | Phoneme and orofacial movement sweeps | • 16 repetitions of each of the 39 English phonemes<br>• 20 repetitions each of various orofacial movements (tongue, lips, jaw, cheeks, larynx, forehead, eyelids) |
| 2 | 2022.04.28 (29) | Initial training data day #1 | • 300 Open WebText training sentences |
| 3 | 2022.05.03 (35) | Individual words from the 50-word set [8] | • 20 repetitions of each of 50 individual words from the Moses et al 2021 vocabulary [8] |
| 4 | 2022.05.05 (36) | Initial training data day #2 | • 380 Open WebText training sentences |
| 5 | 2022.05.17 (49) | Initial training data day #3 | • 440 Open WebText training sentences<br>• 50 Moses sentences used for offline assessment (not training) |
| 6 | 2022.05.19 (51) | Initial training data day #4 | • 200 Open WebText training sentences<br>• 50 Moses sentences used for offline assessment (not training) |
| 7 | 2022.05.24 (56) | Real-time decoding pilot day #1 | • 480 Switchboard training sentences<br>• 75 Moses evaluation sentences |
| 8 | 2022.05.26 (58) | Real-time decoding pilot day #2 | • 480 Switchboard training sentences<br>• 50 Moses evaluation sentences |
| 9 | 2022.06.02 (64) | Real-time decoding pilot day #3 | • 520 Switchboard training sentences<br>• 25 Switchboard evaluation sentences<br>• 50 Moses evaluation sentences |
| 10 | 2022.06.07 (69) | Real-time decoding pilot day #4 | • 480 Switchboard training sentences<br>• 40 Switchboard evaluation sentences<br>• 50 Moses evaluation sentences |
| 11 | 2022.06.14 (76) | Real-time decoding pilot day #5 | • 440 Switchboard training sentences<br>• 40 Switchboard evaluation sentences<br>• 50 Moses evaluation sentences |
| 12 | 2022.06.16 (78) | Real-time decoding pilot day #6 | • 440 Switchboard training sentences<br>• 40 Switchboard evaluation sentences<br>• 50 Moses evaluation sentences |
| 13 | 2022.06.21 (83) | Real-time decoding pilot day #7 | • 400 Switchboard training sentences<br>• 40 Switchboard evaluation sentences<br>• 50 Moses evaluation sentences |
| 14 | 2022.06.23 (85) | Real-time decoding pilot day #8 (silent speaking) | • 440 Switchboard training sentences<br>• 80 Switchboard evaluation sentences<br>• 50 Moses evaluation sentences |
| 15 | 2022.06.28 (90) | Real-time decoding pilot day #9 | • 440 Switchboard training sentences<br>• 80 Switchboard evaluation sentences<br>• 50 Moses evaluation sentences |
| 16 | 2022.07.05 (97) | Real-time decoding pilot day #10 | • 400 Switchboard training sentences<br>• 80 Switchboard evaluation sentences<br>• 50 Moses evaluation sentences |
| 17 | 2022.07.14 (106) | Real-time decoding pilot day #11 | • 440 Switchboard training sentences<br>• 80 Switchboard evaluation sentences<br>• 50 Moses evaluation sentences |
| 18 | 2022.07.21 (113) | Real-time decoding evaluation day (vocal #1) | • 440 Switchboard training sentences<br>• 80 Switchboard evaluation sentences<br>• 50 Moses evaluation sentences |
| 19 | 2022.07.27 (119) | Real-time decoding evaluation day (vocal #2) | • 440 Switchboard training sentences<br>• 80 Switchboard evaluation sentences<br>• 50 Moses evaluation sentences |
| 20 | 2022.07.29 (121) | Real-time decoding evaluation day cut short due to equipment failure (training data only) | • 240 Switchboard training sentences |
| 21 | 2022.08.02 (125) | Real-time decoding evaluation day (vocal #3) | • 440 Switchboard training sentences<br>• 80 Switchboard evaluation sentences<br>• 50 Moses evaluation sentences |
| 22 | 2022.08.11 (134) | Real-time decoding evaluation day (vocal #4) | • 260 Switchboard training sentences<br>• 80 Switchboard evaluation sentences<br>• 50 Moses evaluation sentences |
| 23 | 2022.08.13 (136) | Real-time decoding evaluation day (vocal #5) | • 260 Switchboard training sentences<br>• 80 Switchboard evaluation sentences<br>• 50 Moses evaluation sentences |
| 24 | 2022.08.18 (141) | Real-time decoding evaluation day (silent #1) | • 400 Switchboard training sentences<br>• 80 Switchboard evaluation sentences<br>• 50 Moses evaluation sentences |
| 25 | 2022.08.23 (146) | Real-time decoding evaluation day (silent #2) | • 480 Switchboard training sentences<br>• 80 Switchboard evaluation sentences<br>• 50 Moses evaluation sentences |
| 26 | 2022.08.25 (148) | Real-time decoding evaluation day (silent #3) | • 480 Switchboard training sentences<br>• 80 Switchboard evaluation sentences<br>• 50 Moses evaluation sentences |

### 1.7 Instructed delay tasks

All tasks employed an instructed delay paradigm, with each trial consisting of an instructed delay phase followed by a go phase. For sentence speaking blocks, during the delay period the text of the sentence was displayed on the screen above a red square, providing T12 time to read it and prepare to speak. After the delay period, the red square cue then turned green, and the sentence remained on the screen while T12 attempted to speak it (either aloud or by silently mouthing it, depending on the session). When T12 finished speaking the sentence, she pushed a button held in her lap, which triggered the system to move to the next sentence trial.

For the single phoneme task, each phoneme was cued during the delay period with both text and an audio sample of that phoneme being spoken. Vowels were spoken in isolation and cued with the text of a word containing that vowel, with the vowel capitalized (e.g., strUt for 2). Consonants were all paired with the vowel “ah” following the consonant (denoted “AA” in ARPAbet notation and “A” in IPA). Consonants were paired with a text cue evoking the sound of that consonant (e.g. “kah” or “wah”). For the single word task, where T12 spoke individual words from the 50-word Moses et al. word set [8], each word was cued with text only. For the orofacial movement sweep task, each movement was cued with a text description of that movement (e.g., “Tongue Up”).

We ran a single “diagnostic” block at the beginning of each speech decoding session. Data from the diagnostic block was used to set electrode thresholds and linear regression reference (LRR) coefficients, and was also used to examine the rate of change of neural tuning across days. In this block, T12 spoke individual words from a diagnostic set of 7 words designed to span the space of articulation (with 8 repetitions per word). Words were cued with text only. The word set consisted of the following words: ‘bah’, ‘choice’, ‘day’, ‘kite’, ‘though’, ‘veto’, ‘were’.

Delay period durations were pseudorandomly drawn from an exponential distribution with a task-specific mean (see Table 3); values that fell outside of a specified task-specific range were re-drawn. Go period durations were set fixed to a task-specific value for non-sentences tasks (for the sentence production task, T12 advanced the trial by pressing a button). Finally, in the orofacial movement sweep task, the go period was followed by a short “return” phase where T12 relaxed back to a neutral posture before starting the next trial. In all other tasks, the next trial started immediately after the go period ended.

**Table 3.** Task timing parameters.

| Task | Delay Low | Delay Mean | Delay High | Go Period | Return Period |
| --- | --- | --- | --- | --- | --- |
| Sentences | 4.0 s | 4.5 s | 5.0 s | Variable (button-controlled) | None |
| Phonemes | 1.8 s | 2.3 | 2.8 | 1.7 s | None |
| Orofacial movements | 2.0 s | 2.5 | 3.0 | 1.0 s | 1.0 s |
| Single words (50-word set) | 2.0 s | 2.5 | 3.0 | 2.0 s | None |
| Diagnostic block (7 words) | 1.5 s | 2.0 | 2.5 | 2.0 s | None |

### 1.8 Voiced vs. silent speaking behavior

For most speaking sessions, T12 was instructed to attempt to produce voiced speech in a “typical” manner (i.e., by trying to move all of her articulators and modulate her larynx to pass sound as one would to attempt to speak normally). The acoustic output was largely unintelligible. During the course of the study, T12 reported that due to her reduced breath control abilities, attempting to produce voiced speech was fatiguing. We experimented with different speaking behavior paradigms and found that “mouthing” or silent speaking yielded similar decoding performance to voiced speech while being less fatiguing for T12. For silent speech sessions, we instructed T12 to pretend that she was mouthing the sentence to someone across the room. During this silent speaking behavior, T12 produced no audible sound, but visibly moved her lips, tongue and jaw. Sessions 14, 24, 25 and 26 were all performed with this silent speaking behavior.

### 1.9 Decoder evaluation sessions

Real-time speech decoding was evaluated in sessions 18, 19, 21, 22 and 23 for voiced speaking behavior and sessions 24, 25 and 26 for silent speaking behavior. These were the evaluation sessions reported in Fig. 2 and were conducted with the final version of the real-time decoder and parameters. Previous real-time decoding sessions were pilot sessions used to explore different online approaches and parameters.

Each evaluation session began with a “diagnostic” block as described previously (section 1.7). This block was used to calculate the threshold values and filters for online LRR that would be used for the rest of the session. Then, we collected “open-loop” blocks of sentences (∽6 blocks with 40 sentences per block) during which no decoder was active. We trained the decoder using these blocks of data (combined with data from all past sessions) to obtain a “stage 1” RNN decoder. Next, we collected additional training blocks with real-time feedback, where the decoder output from the stage 1 RNN was displayed in real-time as T12 attempted to speak each sentence. Upon completion of each sentence, T12 would push a button to indicate she was finished and the decoded text was read aloud using Google Cloud’s Text-To-Speech functionality. After the response was voiced by the computer, T12 pushed the button again to continue to the next sentence. Stage 1 real-time decoding was done for 3-6 blocks, with 40 sentences per block. Finally, the RNN was retrained a second-time using all open-loop and stage 1 data (combined with data from all past sessions) to yield a “stage 2” RNN. The stage 2 RNN was then evaluated on the 50 sentences from Moses et al 2021 [8] as well as 80 randomly-selected sentences from the Switchboard corpus, for which the final error rate and word per minute values were reported.

### 1.10 Sentence selection

For the first four initial training data collection sessions, sentences were randomly drawn from the OpenWebText2 corpus [9]. For all subsequent sessions, training sentences were drawn from the Switchboard corpus of telephone conversations between speakers of American English [10]. Sentences were selected by first generating lists of potential sentence segments by automatically splitting the transcription using provided punctuation marks. Sentence segments were then filtered to include only those that expressed a complete meaning, and superfluous starter words (e.g. “and”) were deleted. We also did not include sentences with confusing or distracting meaning, such as violent or offensive topics. Finally, we upsampled sentences with rare phonemes to ensure there was sufficient training data for the RNN to learn these rare phonemes. This resulted in a diverse sample of sentences from spoken conversational contexts.

For each evaluation day, after the final “stage 2” RNN was trained, three evaluation blocks were run. These included one block of the 50 sentences used for evaluation in Moses et al 2021 [8] (these sentences were the same every session), and two blocks of 40 sentences each from Switchboard, selected in the same manner as the training data. The Switchboard sentences were different for each evaluation session. The RNN decoder was never evaluated on a sentence that it had been trained on, and every sentence was unique (except for the “direct comparison” blocks that always used the same 50 sentences from Moses et al 2021). When we retrained the decoder each day before performance evaluation, we retrained it using all previously collected data (from all prior days) except for these direct comparison blocks, in order to prevent the RNN from overfitting to these repeated sentences.

## 2. Neural representation of orofacial movements and speech in orofacial cortex

### 2.1. Tuning heat maps

To generate the neural tuning heat maps shown in Figure 1, we first started with binned threshold crossing spike counts using a –4.5 RMS threshold (20 ms bins). To account for drifts in mean firing rates across the session, the binned threshold crossing rates were mean-subtracted within each block (i.e., for each electrode, its mean firing rate within each block was subtracted from each time step’s binned spike count).

Next, for each trial and electrode, threshold crossing counts were averaged in an 800 ms window (200 to 1000 ms after the go cue). Significance of tuning was then assessed via 1-way ANOVAs applied per electrode, where each ANOVA group corresponded to a different movement condition of a given movement type, and each observation was a scalar average firing rate for a single trial. The movement conditions that were used to assess tuning to each movement type are shown in SFig 3. P-values from each ANOVA were used to define tuning significance (p<1e-5) for the tuning heatmaps (Fig 1f) and mixed tuning counts (SFig 4). To make the statistical power of the “phonemes” and “words” movement types comparable to the orofacial movement types (so that the amount of tuning to phonemes/words can be fairly compared to orofacial movement), we used only the following 6 phoneme or word conditions as opposed to using all 39 phonemes and all 50 words: (B, HH, N, TH, Z, IY) and (am, coming, good, hungry, no, tell). The results appeared robust to the particular set of phonemes/words chosen.

The fraction of variance accounted for by movement tuning on a single electrode was defined as:

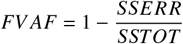

SSTOT is the total sum of squared average firing rates over all trials. For computing SSTOT, squaring was performed after the grand mean across all trials was subtracted from each trial first, so that the overall mean firing rate did not contribute to the variance.

SSERR is the sum of squared prediction errors across all trials. Prediction error was assessed with a cross-validated (5-fold) model which predicts the firing rate of each trial based only on the mean of the condition it belongs to. Condition-specific means were estimated on the training set by taking the sample means across training trials, and then applied to the held-out test set.

If there are large differences in mean firing rate between movement conditions (i.e., strong movement tuning), then SSERR will be small relative to SSTOT. Cross-validation prevents overestimation of tuning due to spurious differences in mean firing rate between conditions that are not stable across folds.

### 2.2. Naive Bayes classification

Offline classification results (reported in Fig 1d,e and SFig 3) were generated using a cross-validated (leave-one-out) Gaussian naive Bayes classifier, following the methods described in [11], using threshold crossing rates computed in a window from 0 to 1000 ms after the go cue (Fig 1d, SFig 3) or a sliding 100 ms window (Fig 1e). We used –4.5 × RMS thresholds for classification. 95% confidence intervals for classification accuracies were computed with bootstrap resampling (10,000 resamples). We chose to use a Gaussian naive Bayes classifier because it is a simple method that performed well enough to demonstrate the existence of strong neural tuning - it is likely that more advanced methods could improve classification accuracy further.

### 2.3. Preserved articulatory representation of phonemes

#### 2.3.1. Electromagnetic articulography (EMA) representations

Electromagentic articulography (EMA) and corresponding audio data was taken from the publicly available “USC-Timit” dataset [12] and the “Haskins Production Rate Comparison” dataset [13], which both contain phoneme labels (the beginning and end of each phoneme is marked for each sentence). EMA data was collected with markers placed on the lip, tongue and jaw. We used only the X and Y positions of each marker (sagittal plane position), yielding an articulatory time series with dimensionality ranging from 12 (6 markers) to 16 (8 markers) depending on the subject.

To compute an EMA representation of each phoneme, we averaged over all EMA marker position data recorded at time points that were labeled as belonging to that phoneme, yielding a single average articulator position vector for each phoneme. Finally, a binary voicing indicator variable was added to the EMA representation of each phoneme. This variable was set to 1 for voiced phonemes (e.g., ‘z’) and 0 for unvoiced phonemes (e.g., ‘s’). Since EMA does not measure voicing, this is a simple way to include some voicing information that would otherwise be omitted from the EMA representation.

EMA data from USC-Timit subject ‘M1’ was used for all Figure 3 analyses. SFig 8 shows data from all 12 subjects.

#### 2.3.2. Saliency vector representation

Saliency vectors, which quantify the neural vectors which maximally excite each of the decoder’s phoneme outputs, were generated by computing RNN logit gradients with respect to the input features. We used an RNN trained on all voiced speech days and then used the first 30 trials from each day’s test set to compute the gradients.

For each day’s data, we run the RNN over each sample

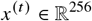

for five time steps (first initializing the hidden state to zeros) - this allows the network some time to integrate information about the specific feature vector. This yields a phoneme probability output for each time step

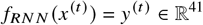

For each time step *t*, we then calculate the Jacobian matrix *j*, which contains entries corresponding to first-order partial derivatives of each logit output with respect to each channel, i.e.

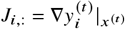

This gradient records how small changes in each channel’s activity influences the probability of class *i*. We can calculate the Jacobian for all phonemes and timesteps, resulting in a time x channels x phonemes matrix M where

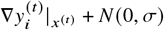

contains the Jacobian at timestep *t*. We then average across the time dimension to obtain an integrated estimate of how each channel’s activity influences different phoneme class probabilities.

To compute the gradients, we used SmoothGrad [14], a method for denoising saliency maps by computing gradients over an input with multiple noise perturbations, i.e.

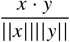

The resulting saliency map estimates are then averaged together. We use n = 20 perturbations and noise level = 10 (relative to the overall range of firing rates after capping outliers). This extra step contributed a small but consistent improvement in similarity matrix correlations with the EMA data.

#### 2.3.3. Similarity matrices

Similarity matrices in Figure 3a and 3d were computed using cosine similarity. That is, for each pair (*x, y*) of RNN saliency or EMA vectors, similarity was defined as

This equation computes the cosine of the angle between *x* and *y*. Before computing cosine similarity, the vectors were first centered by subtracting the mean across all consonant (Figure 3a) or all vowel vectors (Figure 3d).

#### 2.3.4. Correlation between articulatory and neural representations

Figure 3b and 3e show the correlation between the neural and EMA phoneme representations, which was computed across all phonemes and dimensions after a cross-validated Procrustes alignment of the saliency vectors to the EMA vectors. In SFig 8, this same method was used to compute the correlation between neural representations and articulatory representations that were estimated in different ways. In SFig 8c, correlation values were computed between consonants and vowels separately and then averaged together to produce a single value.

First, neural vectors were concatenated into a 256 × 39 matrix **X** (256 neural features x 39 phonemes) and articulatory vectors were concatenated into a N x 39 matrix **Y** (N articulatory dimensions x 39 phonemes). **X** and **Y** were then reduced to D dimensions using PCA applied across the rows, yielding an D x 39 matrix 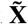 and an D x 39 matrix 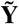. D was set to 12 if N>=12, otherwise D was set equal to N.

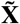 was then aligned to 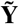 using a cross-validated orthogonal rotation (Procrustes analysis) using leave-one-out cross-validation. Specifically, for each vector *x*_*i*_ in 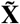, Procrustes analysis was applied to align all other vectors {*x*_1_, …*x*_*i−*1_, *x*_*i*+1_, …*x*_*n*_} to the matching vectors {*y*_1_, …*y*_*i−*1_, *y*_*i*+1_, …*y*_*n*_}, yielding an orthogonal rotation **R. R** was then applied to *x*_*i*_ to yield 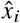. Orthogonality enforces that the rotation be rigid, so that the underlying structure in the data is preserved.

Finally, all 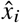 vectors were concatenated into an (8×39) × 1 vector 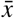 and all 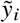 vectors were concate-nated into an (8×39) x 1 vector 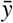. The Pearson correlation coefficient was then calculated between 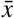 and 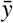 using consonant entries only (Figure 3b) or vowel entries (Figure 3e).

As a control, this same procedure was repeated 10,000 times but with the columns of **X** shuffled into a random order, which allows estimation of what the correlation could be expected to be ‘by chance’ if each phoneme’s vector was random but drawn from the same distribution. Note that the cross-validation procedure causes the chance distribution to be centered at 0 (otherwise it would be biased upwards as Procrustes would overfit and align noise). The true correlation is far greater than any of the 10,000 shuffle results, indicating statistical significance (p<1e-4).

#### 2.3.5. Low-dimensional visualization of phoneme geometry

To make the plots in Figure 3c and 3f, the neural saliency vectors were first aligned to the EMA vectors using cross-validated Procustes analysis, as described in the above section. After alignment, the top two dimensions were plotted; for vowels, these two dimensions were rotated and flipped within the plane in order to highlight the classic (front vs. back) and (high vs. low) structure.

#### 2.3.6. Place and formant representation

In SFig 8, we tested a different method for quantifying the articulatory representation of phonemes based on classical ways of understanding consonants and vowels. We represented each consonant based on its place of articulation (labial, dental, alveolar, post-alveolar, velar, glottal) by using a 24 × 6 “place code” matrix. In this matrix, each row is a consonant and each column corresponds to one of the six places of articulation. For each consonant, we placed a “1” in the column corresponding to its place of articulation and a “0” in all other columns. We represented vowels using a 15 × 2 “formant” matrix.

In this matrix, each row is a vowel, and the two columns correspond to the logarithm of the first and second formant frequencies [15].

#### 2.3.7. Decoder labeling representation

In SFig 8, we tested different methods for quantifying the neural representation of phonemes. One of these was the “Decoder Labels” method, which was based on identifying time steps that belonged to each phoneme using the RNN decoder output, and then averaging across these time steps to find a neural representation vector for each phoneme.

To accomplish this, we used the segmentation algorithm from [16]. This algorithm estimates a time window during each phoneme occurs, given the outputs of a decoder trained with the CTC loss function [17]. For this approach, we trained a bidirectional RNN offline, because previous studies have found that bidirectional RNN strained with the CTC loss produces better alignment than unidirectional RNNs [18, 19]. Because RNN models trained with the CTC loss tend to delay their predictions [18, 19], we extended the start time of each phoneme’s neural activity segment by one kernel size (defined in Table 5). Once the time windows for each phoneme were identified, we simply averaged over all time steps to generate a single average neural vector for each phoneme. For offline RNN training, we followed the procedure described below in section 7.

#### 2.3.8. Single phoneme task representation

In SFig 8, we tested different methods for quantifying the neural representation of phonemes. One of these was the “Single Phoneme Task” method, which was based on the instructed delay task shown in Fig. 1. In this task, T12 spoke individual vowels or consonants paired with the same vowel “AH”. To compute the neural representation of each phoneme, we simply averaged the threshold crossing rates within a 0 to 1000 ms window after the go cue for each trial belonging to that phoneme (threshold = -4.5 RMS).

### 2.4 Neural correlation across days

To compute how the neural representation of speech was correlated between pairs of days (Fig. 4d), we used data from a “diagnostic block” collected at the beginning of each day. During this block, T12 completed an instructed delay task where she attempted to speak individual words from a set of 7 words designed to span the space of articulation (8 repetitions per word). The word set consisted of the following words: ‘bah’, ‘choice’,’day’,’kite’,’though’,’veto’, and ‘were’. We also included a condition where T12 was instructed to rest silently (‘do nothing’).

First, threshold crossing rates for each trial were averaged between a 100 to 600 ms window after the go cue to yield a single firing rate vector for each trial (of length 128). Then, “pseudo-trial” vectors were created by concatenating together a single firing rate vector from each condition, resulting in pseudo-trial vectors of length 128*8=1024. The result of this step is a set of eight vectors {*v*_1_, *v*_2_,, *v*_8_}, one vector for each of the eight repetitions of all conditions. When assessing the similarity between any two days, we then have two sets of vectors to consider: {*v*_1_, *v*_2_, …, *v*_8_} and {*u*_1_, *u*_2_,, *u*_8_}. Consider each of these vectors as a random draw from a day-specific distribution (let us denote the two distributions as *V* and *U*). To quantify similarity, we estimated the correlation between the means of *V* and *U* (note that the means themselves are also vectors). The quantity of interest here is the mean because this represents the average firing rates observed for each condition (i.e., the neural representation of each word). To estimate the correlation between the means of *V* and *U*, we used a cross-validated measure of correlation that reduces the impact of noise. See our prior work [11] and accompanying code repository https://github.com/fwillett/cvVectorStats for more details about this method. Importantly, this cross-validated method is different from simply correlating 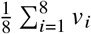 and 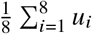 which would underestimate the true correlation due to noise that causes the estimated means to appear more dissimilar than they really are. For example, even if *V* and *U* have identical means, noise in *v*_*i*_ and *u*_*i*_ would always cause the estimated correlation to be less than 1 when correlating 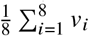and 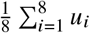 together.

To make the plot in Fig. 4d, we included all pairings of the following 12 days on which a diagnostic block of attempted vocal speaking was collected: 2022.06.16, 2022.06.21, 2022.06.28, 2022.07.05, 2022.07.07, 2022.07.14, 2022.07.21, 2022.07.27, 2022.07.29, 2022.08.02, 2022.08.11, 2022.08.13.

## 3. Phoneme transcription and labelling

Each sentence prompt was transcribed into a sequence of phonemes using the CMU Pronouncing Dictionary (http://www.speech.cs.cmu.edu/cgi-bin/cmudict) and the g2p software package [20]. The CMU Dictionary uses 39 phonemes, each denoted using the ARPAbet symbol set developed for speech recognition (see Table 4 for the correspondence between IPA notation and ARPAbet notation, and https://en.wikipedia.org/wiki/ARPABET). Note that we did not incorporate the stress labeling given by the CMU dictionary for vowels (i.e., we labeled each vowel in the same way regardless of how it is stressed in the word).

We automatically added a “silence” phoneme to the end of each word in order to denote the separation between words. We reasoned that this gives the RNN decoder the ability to communicate demarcations between words to the language model, and we found that it improved performance. Note that the silence token is not necessarily intended to model literal silence, as the participant may or may not be silent between each word (although she appeared to take brief pauses between many of the words, as can be seen in SVideo 1).

## 4. Decoder performance metrics

### 4.1. Word and phoneme error rates

We evaluate both phoneme error rate and word error rate. Phoneme error rate was defined as the edit distance between the decoded sequence of phonemes and the prompt sentence phoneme transcription (i.e., the number of insertions, deletions or substitutions required to make the sequence of phonemes match exactly). Similarly, word error rate was the edit distance defined over sequences of words.

Note that the reported error rates are the combined result of many independent sentences. To combine data across multiple sentences, we summed the number of errors across all sentences and divided this by the total number of phonemes/words across all sentences (as opposed to computing error rate percentages first for each sentences and then averaging the percentages). The helps prevent very short sentences from overly influencing the result.

Confidence intervals for error rates were computed via bootstrap resampling over individual trials and then re-computing the error rates over the resampled distribution (10,000 resamples).

### 4.2. Words per minute

Words per minute was defined as the number of words spoken divided by the total amount of speaking time. Speaking time for each trial was defined as the time from which the cue turned green to when the participant pushed the button to signal she had completed saying the prompted sentence.

Confidence intervals for words per minute were computed via bootstrap resampling over individual trials and then re-computing the speaking rate over the resampled distribution (10,000 resamples).

## 5. RNN architecture

We used a 5 layer, stacked gated recurrent unit RNN [21] to convert T12’s neural activity into a time series of phoneme probabilities. The RNN ran at a 4-bin frequency (20 ms bins), outputting a phoneme probability vector every 80 ms. A 14-bin window of neural activity was stacked together and fed as input to the RNN at each 80 ms cycle (in other words: kernel size = 14, stride = 4). See SFig 5 for parameter sweeps that justify these and other architecture choices.

**Table 4.**
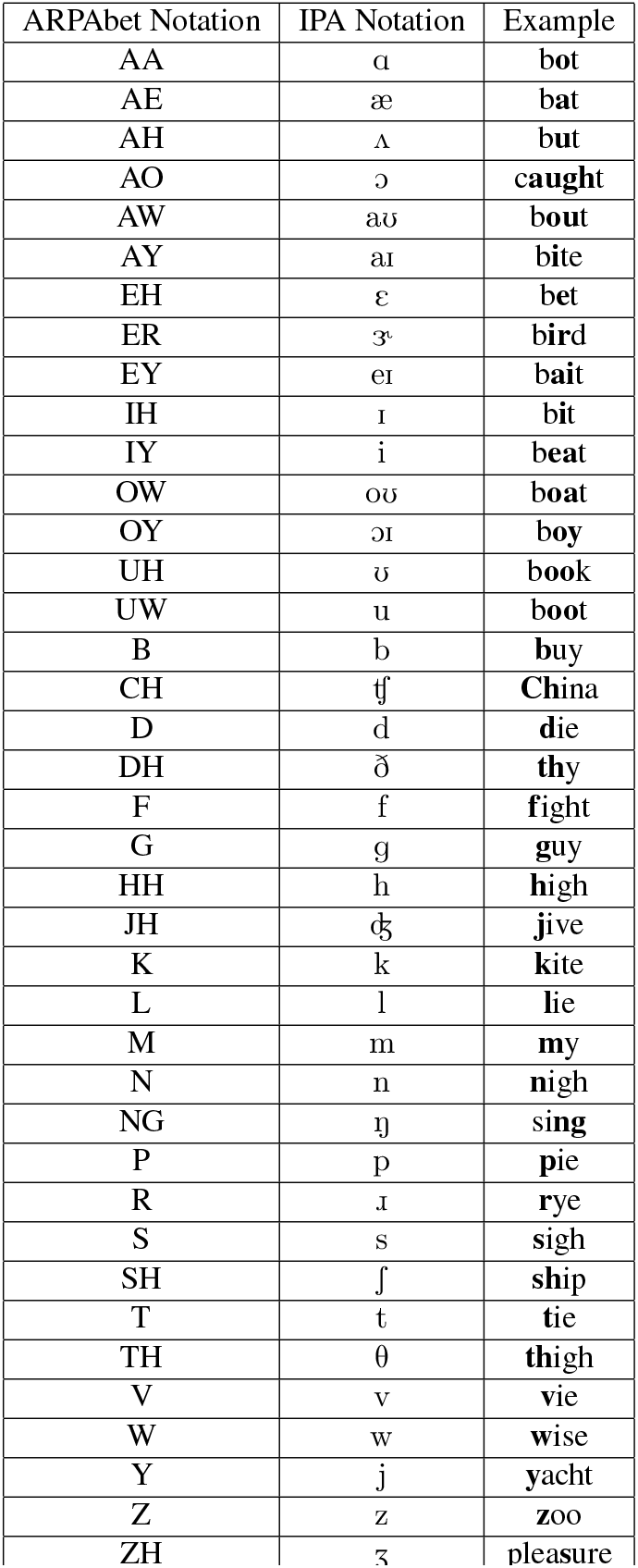
ARPAbet and IPA correspondence.

| ARPAbet Notation | IPA Notation | Example |
| --- | --- | --- |
| AA | ɑ | bot |
| AE | æ | bat |
| AH | ʌ | but |
| AO | ɔ | caught |
| AW | aʊ | bout |
| AY | aɪ | bite |
| EH | ɛ | bet |
| ER | ɜ˞ | bird |
| EY | eɪ | bait |
| IH | ɪ | bit |
| IY | i | beat |
| OW | oʊ | boat |
| OY | ɔɪ | boy |
| UH | ʊ | book |
| UW | u | boot |
| B | b | buy |
| CH | tʃ | China |
| D | d | die |
| DH | ð | thy |
| F | f | fight |
| G | g | guy |
| HH | h | high |
| JH | dʒ | jive |
| K | k | kite |
| L | l | lie |
| M | m | my |
| N | n | nigh |
| NG | ŋ | sing |
| P | p | pie |
| R | ɹ | rye |
| S | s | sigh |
| SH | ʃ | ship |
| T | t | tie |
| TH | θ | thigh |
| V | v | vie |
| W | w | wise |
| Y | j | yacht |
| Z | z | zoo |
| ZH | ʒ | pleasure |

### 5.1. Feature pre-processing and day-specific input layers

Threshold crossing rates and spike band power features were pre-processed by binning into 20 ms time steps, “z-scoring” (mean-subtracted and divided by the standard deviation), causally smoothed by convolving with a Gaussian kernel (sd = 40ms) that was delayed by 160ms, concatenated into 256 × 1 vector, and then transformed using a day-specific input layer. Z-scoring was performed using block-specific means and standard deviations (to account for non-stationarities in the features that accrue over time across blocks). Using day-specific input layers outperformed the alternative of a shared input layer across all days (SFig 5B).

The day-specific input layers consisted of an affine transformation applied to the feature vector followed by a softsign activation function:

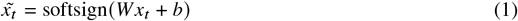

Here, 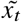 is the day-transformed input vector at time step *t, W*_*i*_ is a 256 × 256 matrix and *b*_*i*_ is a 256 × 1 bias vector for day *i*, and the softsign function is applied element-wise to the resultant vector (where softsign 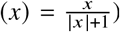.*W*_*i*_ and *b*_*i*_ were optimized simultaneously along with all other RNN parameters. During training, dropout was applied both prior to and after the softsign.

### 5.2. Rolling z-scoring

During online evaluation, we used a rolling estimate of the mean and standard deviation of each feature to perform z-scoring. This helps account for neural non-stationarities that accrue across time, and substantially outperforms the alternative of using the prior block’s means and standard deviations (SFig 5A).

For the first ten sentences of a new block, we used a weighted average of the prior block’s mean estimate and the mean of whatever sentences were collected so far in the current block:

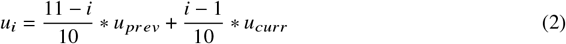

Here, *u*_*i*_ is the mean used to z-score sentence *i, u* _*prev*_ is the prior block’s mean estimate, and *u*_*curr*_ is the mean across all sentences collected so far in the current block. After ten sentences had been collected, we stopped incorporating the prior block’s mean and simply took the mean across the most recent min(20, *N*) sentences, where *N* is the number of sentences collected so far in the current block. The standard deviation was updated in the same way as the mean.

## 6. RNN training overview

### 6.1. Connectionist temporal classification (CTC) loss

Due to T12’s inability to produce intelligible speech, we had no ground truth labels of what phonemes were being spoken at each time step. The lack of ground truth labels makes it difficult to apply simple supervised training techniques to train the RNN. To get around this problem, we used the Connectionist Temporal Classification (CTC) loss function, which can train neural networks to output a sequence of symbols (in this case, phonemes) given unlabeled time series input [17]. Using the CTC loss function results in an RNN that is trained to output a time series of phoneme probabilities (with an extra “blank” token probability). A language model can then be used to infer a sequence of underlying words from these probabilities, or phonemes can be decoded from these probabilities simply by emitting the phoneme of maximum probability at each time step (while taking care to omit repeats and time steps where “blank” is the maximum probability).

### 6.2. Artificial noise

We added two types of artificial noise to the neural features to regularize the RNN. First, we added white noise directly to the input feature vectors at each time step. Adding white noise to the inputs asks the RNN to map clouds of similar inputs to the same output, improving generalization. We also added artificial constant offsets to the means of the neural features, to make the RNN more robust to non-stationarities in the neural data. Drifts in the baseline firing rates that accrue over time has been an important problem for intracortical BCIs [22, 23, 24]. The constant offset values were randomly chosen on each minibatch and were constant across all time steps in the minibatch, but unique to each feature.

The two above-mentioned types of noise (white noise and constant offset noise) were combined together to transform the input vector in the following way:

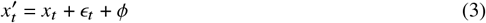

Here, 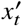 are the neural features with noise added, *x*_*t*_ are the original neural features, ϵ_*t*_ is a white noise vector unique to each time step, and *ϕ* is a constant offset vector.

### 6.3. Supervised training

The RNN was implemented with TensorFlow 2 and trained using stochastic gradient descent (ADAM; *β*_1_ = 0.9, *β*_2_ = 0.999, *ϵ* = 0.1) for 10,000 minibatches (batch size = 64). The learning rate was decayed linearly from 0.02 to 0.0 across the 10,000 minibatches. We applied dropout and L2 weight regularization during training to improve generalization. See Table 5 for a list of RNN hyperparameters.

**Table 5.** Architecture, training and regularization parameters for RNN Model.

| RNN Parameter | Description | Value |
| --- | --- | --- |
| nUnits | Number of units in each GRU layer | 512 |
| nLayers | Number of GRU layers | 5 |
| Kernel Size | Number of input feature time bins stacked together as a single input for the RNN | 14 |
| Stride | Describes how many time bins the RNN skips forward every step | 4 |
| L2 | L2 regularization cost | 1e-0.5 |
| Dropout | Probability of dropout during training | 0.4 |
| WhiteNoiseSD | Standard deviation of white noise added to input data for regularization | 1.0 |
| constantOffsetSD | Standard deviation of constant offset noise added to input data to improve robustness against non-stationary feature means | 0.2 |
| Batch Size | Number of sentences included in each mini-batch | 64 |
| Learning Rate | Linearly decaying learning rate | 0.02 to 0.0 |
| $\beta_1$ | ADAM stochastic gradient descent parameter | 0.9 |
| $\beta_2$ | ADAM stochastic gradient descent parameter | 0.999 |
| $\epsilon$ | ADAM stochastic gradient descent parameter | 0.1 |

## 7. Offline performance sweeps

### 7.1. Overview

To determine the effect of different design choices made for the RNN architecture, and to understand the impact of data quantity and channel count, we performed several performance sweeps offline (results from this are shown in SFig 5, SFig 7 and Fig 4). Unless otherwise specified, we trained 10 seeds of an RNN model for each variation of parameters and performed inference on a standardized set of held-out test data. To define the training/held-out set, we took 40 sentences at random from each day as the held-out set and used the remaining sentences as offline training data, drawing from all open-loop/stage 1 sentences on each day (but excluding stage 2 evaluation data). We used the original language model that was run online (as opposed to the improved version reported in Table 1).

We ran parameter sweeps for the GRU-RNN architecture choices including number of units, number of layers, kernel size and stride (SFig 5e-h). A comparison between a shared input network and unique input network per session was also run (SFig 5b). Furthermore, performance when using different kinds of features was also compared in this manner (SFig 5c-d), including four different threshold crossing thresholds (−3.5,-4.5,-5.5,-6.5), spike band power, area 44 vs. area 6v features, and the mel-frequency cepstral coefficients (MFCCs) of the participants’ recorded audio during attempted speaking sessions. MFCCs were computed using 40 ms window (using MATLAB 2020a’s “mfcc” function).

### 7.2. Effect of channel count on performance

To determine the effect of channel count on decoding performance (Fig 4b), 100 seeds of each multiple of 10 number of channels up to the full 128 channels was run. For each of the 100 seeds for each channel count, channels were randomly selected without replacement. To predict performance for higher number of channels past 128, a least squares linear regression was fit to the log-log relationship of the number of channels vs. error rate.

### 7.3. Amount of training data

To plot the number of days of training data versus performance (SFig 5i), RNNs were trained for each of the 5 vocal speaking evaluation days separately, and for each number of training data days going consecutively in reverse until all previous days were used in training. Performance was assessed only on the given evaluation day. Word error rates were then averaged over all evaluation days to produce a single (# of days) vs. (word error rate) curve.

We also tested whether or not it was necessary to retrain the RNN decoder on each new performance evaluation day using hundreds of new sentences collected on that day, or whether fewer (or no) new sentences might have also yielded good performance, which would be a more realistic use case (Fig 4c). For this analysis, models were trained on the five attempted speech evaluation sessions (sessions 18,19,21,22,23) using reduced subsets of sentences from the given evaluation day (while still using all historical data). The input layer for each given evaluation day was also tied to be the same as the most recent historical day, in order to prevent overfitting when using a small number of training sentences. Once trained, RNNs were evaluated on the same set of “stage 2” online evaluation sentences used to report performance in Figure 2.

### 7.4. Language model vocabulary size sweep

To test how the number of words in language model (LM) affects the decoding accuracy, we built different 3-gram LMs with various vocabulary sizes. These LMs were built following the same procedure as in Section 8, but with vocabulary sizes varying from 50 to 140,000. We started with the 50 words from [8], and gradually added words until the vocabulary size reached 140,000. The added words were chosen from the LM training corpus. Words were added in the order of their frequencies in the training corpus. When the vocabulary size became greater then 4500, we pruned the LM with threshold 1*e −* 9. To measure the WER, we ran the LM decoders on the CTC probabilities output by RNN from the 8 real-time speech decoding sessions (18, 19, 21, 22, 23, 24, 25, and 26).

## 8. Language model

### 8.1. Overview

We used a n-gram language model (LM) to decode word sequences from RNN outputs for real-time decoding and offline analyses. Here, we give an overview of the major steps involved. The n-gram LM was created with Kaldi [25] using OpenWebText2 corpus [9]. We first preprocessed the text corpus to only include English letters and limited punctuation marks. Then we used Kaldi to construct a n-gram LM, using either the CMU Pronunciation Dictionary ^1^ (125k words) or the 50 words from [8]. The LM was represented in the form of a weighted finite-state transducer [26] which can be used to translate the a sequence of CTC labels into candidate sentences.

### 8.2. OpenWebText2 preprocessing

Our n-gram LM was created using samples from OpenWebText2 [9]. OpenWebText2 is a text corpus covering all Reddit submissions from 2005 up until April 2020. We downloaded the entire corpus and randomly sampled 95% as a training corpus. We preprocessed the training corpus to include only English letters and 4 punctuation marks (period, comma, apostrophe, and question mark). The preprocessed corpus was then split into sentences and converted to upper case (yielding a total of 634M sentences with 99B words).

### 8.3. Constructing the n-gram language model

We used publicly available scripts ^2^ as a starting point for constructing our n-gram LM. The script first uses SRILM [27] to count the frequencies of n-grams (unigram, bi-gram, and 3-gram, etc.) in the training corpus. We used the Good-Turing discounting method [28] to improve probability estimation of unseen or rare word combinations. For words that are not in the pronunciation dictionary, they are mapped to a special token <UNK>. When using the CMU Pronunciation Dictionary, the resutling LM is too large to fit into the main memory of the Ubuntu computer used for real-time inference. We pruned the resulting n-gram LM using SRILM, which removes n-grams that causes the perplexity of the LM to increase by less than a threshold. The LM built with the 50 words from [8] is not pruned. For online real-time decoding, we used a 3-gram LM pruned with threshold 1*e −* 9. For offline analyses, we used a 5-gram LM pruned with threshold 4*e −* 11.

The n-gram LM was then converted to a weighted finite-state transducer (WFST) [26]. A WFST is a finite-state acceptor in which each transition has an input symbol, an output symbol and a weight. A path through the WFST takes a sequence of input symbols and emits a sequence of output symbols. We followed the recipe in [29] to construct our WFST search graph:

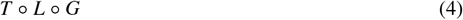

Here, ∘ denotes composition.

G is the grammar WFST that encodes legal sequences of words and their probabilities based on the n-gram LM.

L is the lexicon WFST that encodes what phonemes are contained in each legal word. Note that a “silence” phoneme was added to the end of each word (see section 3). We did an offline sweep of the “silence” phoneme probability and found 0.9 to be optimal.

Finally, T is the token WFST that maps a sequence of RNN output labels to a single phoneme. In our case, T contains all the individual phonemes plus the CTC blank symbol. For more details about how the three WFSTs were composed, refer to [29].

### 8.4. Inference with the n-gram language model

We used the LM decoder implementation in We Net [30] for efficient real-time inference. We Net is a wrapper around Kaldi to simplify the implementation of a real-time LM decoder. The LM decoder runs an approximate Viterbi search (beam search) algorithm on the WFST search graph to find the most likely sequences of words. The WFST search graph encodes the mapping from a sequence of CTC labels emitted by RNN to a sequence of words. During inference, the beam search combines information from the WFST (state transition probabilities) and information from the RNN decoder about which CTC labels are likely occurring at each moment in time. We do not normalize the CTC label probabilities as in [29].

The decoding parameters for beam search in defined in Table 6. The beam search runs every 80ms, after the RNN emits CTC label probabilities. On average, each beam search step took less than 1ms to complete.

### 8.5. Offline language model optimization

After data collection was completed, we further optimized the LM and found that online decoding WER could have improved by 6.4% with an improved LM architecture (Table 1 in main text). To improve the LM, we used a 5-gram LM instead of 3-gram LM and employed a 2-pass decoding strategy. The first pass of the 2-pass decoder is the same as the 1-pass decoder described above. But instead of outputting a decoded sentence, it outputs a word lattice [31, 32]. A word lattice is a directed graph where each node is a word and the an edge between nodes encodes the transition probability between words. It is a efficient representation to encode possible word sequences. The second pass of the 2-pass decoder uses a unpruned n-gram LM to rescore the word lattice. Rescoring replaces the original LM score with a more accurate score from the unpruned LM. After rescoring, we pick the best path through the word lattice as decoding output.

Finally, we found that using a transformer LM [33] to rescore the candidate sentences in an third pass could further improve decoding accuracy. Transformer LMs have been the state of the art in many natural language tasks in recent years [34, 35]. Compared to an n-gram LM which models a limited context (e.g., 3 words for a trigram model), a transformer LM can model much longer contexts (e.g., 1024 words). Training a transformer LM requires a significant amount of computation resources. We used the publicly available pre-trained OPT LM [36]. We used the largest OPT LM (6.7B parameters) that can fit into one NVIDIA A100 40GB GPU. The OPT LM was used to rescore the n-best outputs from a 2-pass decoder.

The 2-pass decoder first outputs at most n sentences with the highest decoding scores. The decoding score of a sentence was defined as follows:

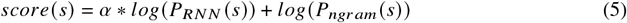

output by the RNN.

Here P _*RNN*_ (*s*) is the sentence *s*’s corresponding CTC label sequence probability *P*_*ngram*_ is the sentence *s*’s probability estimated by the n-gram LM. α is the *acoustic scale* defined in Table 6.

We then used OPT to evaluate the probability of each sentence in the n-best list and linearly interpolate with the n-gram LM’s probability. The new score function was defined as follows:

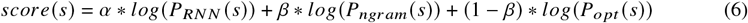

Here *P*_*optt*_ (*s*) is sentence *s*’s probability estimation from OPT LM. *β* is the *lm weight* defined in Table 6. The top scored sentence is the final decoding output.

Finally, we found that decoding accuracy was improved by dividing the CTC blank label probability by a constant value [37], which adds a cost for not outputting any labels.

All LM decoding parameters are optimized via grid search on a validation data set (session 7-16).

**Table 6.**
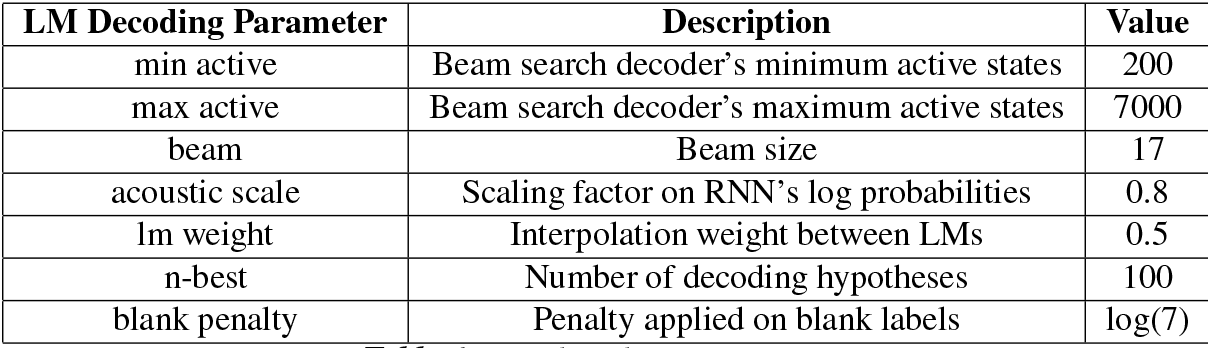
LM decoding parameters.

## Footnotes

1 http://www.speech.cs.cmu.edu/cgi-bin/cmudict

2 https://github.com/thu-spmi/CAT/blob/v1/egs/libri

